# A panoramic view of the expression and function of the Doublesex/DMRT gene family in *C. elegans*

**DOI:** 10.64898/2026.01.01.697290

**Authors:** Chen Wang, Yehuda Salzberg, Meital Oren-Suissa, Oliver Hobert

**Affiliations:** Department of Biological Sciences, Howard Hughes Medical Institute, Columbia University, New York, New York, USA; Weizmann Institute of Science, Department of Neurobiology, Rehovot, Israel

## Abstract

Throughout the animal kingdom, sex determination and sexual differentiation are orchestrated by a strikingly diverse set of regulatory factors. The only type of molecules consistently deployed during sexual differentiation are members of the Doublesex/Mab-3-related transcription factor (DMRT) family. Although each animal genome codes for a multitude of DMRT family members, in no species has the full array of DMRT genes been comprehensively analyzed across the entire animal, in all sexes and throughout development. Hence, the extent of deployment of DMRT genes in sexual differentiation remains unknown. We describe here the first genome- and nervous system-wide expression and functional analysis of all members of the DMRT gene family. Leveraging genome-engineered reporter alleles of all ten DMRT genes of the nematode *Caenorhabditis elegans*, we find that six DMRTs display sexually dimorphic expression in somatic and/or reproductive tissues, including in cell and tissue types not previously known to be sexually dimorphic. In the nervous system, DMRT protein expression covers many, though not all, known sexually dimorphic neuron types. Analyses of DMRT null mutant alleles reveal a suite of neuronal differentiation defects, ranging from altered neurotransmitter identities and switched neuropeptide signatures to impaired glia-to-neuron transdifferentiation. Several DMRT proteins do not exhibit sexually dimorphic expression, indicating roles beyond sexual differentiation. Similar comprehensive analyses of DMRT genes in other organisms may help to better understand the extent and regulation of sex-specific cellular differentiation programs.

## INTRODUCTION

Within most animal species, members of opposing sexes display striking phenotypic differences. Such differences are not restricted to reproductive organs. For example, across mammals, including in humans, sex-biased gene expression is observed in all major somatic organ types, including the brain^1–3^. Sex differences throughout all somatic organ systems are also evident in invertebrate species, with the most extensively studied examples being the fruit fly *Drosophila melanogaster* and the nematode *Caenorhabditis elegans*. Apart from gonadal structures, the best studied sex differences in these model systems lie in the nervous system, ranging from disparities in the sizes of specific brain regions, sex-specific patterns of cell death, connectivity among brain regions and individual cells, to differential gene expression^3–6^.

The genetic regulatory architecture that induces sex-specific differentiation programs throughout the animal is strikingly diverse. In vertebrates, gonadal sex hormones play an important role in instructing sexual differentiation across several somatic cell types^7^. In contrast, in invertebrates, cell-autonomous, sex chromosome-based differences appear to constitute the main drivers of sexual differentiation ^7^. In spite of these apparent differences, one family of regulatory factors is consistently deployed to control sex-specific differentiation programs across the animal kingdom. These regulatory factors are members of the <u>D</u>oublesex and <u>M</u>AB-3-<u>r</u>elated <u>t</u>ranscription factor (DMRT) family, characterized by the presence of a DNA binding “Doublesex/MAB-3” (DM) domain^8–12^. Unlike most other transcription factor families involved in developmental patterning (e.g. homeodomains, Zn fingers, bHLH), DMRT proteins are a metazoan innovation^13^. In vertebrates, the DMRT protein DMRT1 controls gonad differentiation, while in insects, the DMRT protein Doublesex acts as a master regulator of sexual differentiation throughout the entire animal ^8,9^. In *C. elegans*, several DMRT genes are known to control various sexually dimorphic neuronal and non-neuronal features of the animal ^4,8,9,11,14–26^. Even in very basal metazoans, such as cnidarians, DMRT genes are not only known to exist, but are expressed in a sex-specific manner^27^.

While some DMRT proteins have been well studied in the context of sexual differentiation in several animal species, many others have not, particularly outside the gonad. For example, the mouse genome encodes seven DMRT genes, several of which implicated in embryonic brain development, yet only two have been examined for potentially sexually dimorphic function in the brain during sexual differentiation ^10,11,28,29^. Similarly, *in Drosophila*, the Doublesex gene has been extensively studied, but its three DMRT paralogs *dmrt99B, dmrt93B*, and *dmrt11E* have received very little attention ^30^. This lack of knowledge leaves room for the attractive speculation that many and perhaps all members of this gene family operate in sexual differentiation programs of each of these species.

We sought to address the depth by which DMRT proteins contribute to sexual differentiation, using *C. elegans* as a model system. The genome of the nematode *C. elegans* was previously suggested to encode 11 DMRT genes, even more than vertebrates ^12,13,31,32^ (we correct here the number to 10). The founding member, *mab-3*, was initially identified through male patterning defects ^16,33^. Another male tail patterning mutant, *mab-23*, was subsequently recognized to also be a member of the DMRT family ^34^. Ensuing reverse genetic analysis of *dmd-3, dmd-4, dmd-5*, and *dmd-11* also revealed sexually dimorphic functions in skin and neuronal patterning ^18,19,21,35,36^. However, other than the DMD-4 protein, the expression of none of these DMRT proteins has been comprehensively analyzed throughout the entire animal.

Here, we present the first systematic analysis of the entire family of *C. elegans* DMRT proteins, with the overall goal of uncovering the breadth of their influence on sexual differentiation. We specifically set out to address several key questions: Do DMRT proteins mark all sexually dimorphic cells in the animal? Does DMRT expression reveal dimorphisms not previously uncovered in the extensive anatomical analysis of both *C. elegans* sexes? To what extent do DMRT proteins shape sexual dimorphisms in sex-shared organs and/or cells? Is there a common theme in how DMRT genes contribute to cell differentiation, particularly within the nervous system?

## RESULTS

### Nematode DMRT family members

Based on the most recent genome annotation and transcript evidence, the *C. elegans* genome contains a total of ten DMRT genes, one less than previously described ^31^, due to an initially aberrant split of what is now a single locus, *dmd-10*, into two genes (**Fig. 1A-C**). The DMRT complement of *C. elegans* is larger than the seven DMRT genes present in humans and four in *Drosophila* ^11,31,32^. In spite of this larger DMRT complement, *C. elegans* has lost one of the ancestral and deeply conserved set of three DMRT genes ^32^. Within this ancestral set (termed “group 1”), *C. elegans* has a single ortholog of the *DMRT4/5* gene (*dmd-5*) and of the *DMRT93B* gene (*dmd-4*), but it has lost a *DMRT2* representative. The remaining eight *C. elegans* DMRT genes are nematode-specific expansions that fall into two groups (“group 2” and “group 3”), neither of which is chromosomally clustered (**Fig. 1A**). Group 2 is defined by the closely related *dmd-6, dmd-7*, and *dmd-9* genes. Group 3 is defined by the related *mab-3, mab-23, dmd-3, dmd-8*, and *dmd-10* genes. Four of those five group 3 genes encode two adjacent DM domains each (**Fig. 1A**), a feature not shared by DMRT genes in other invertebrates or vertebrates.

**Figure 1.**
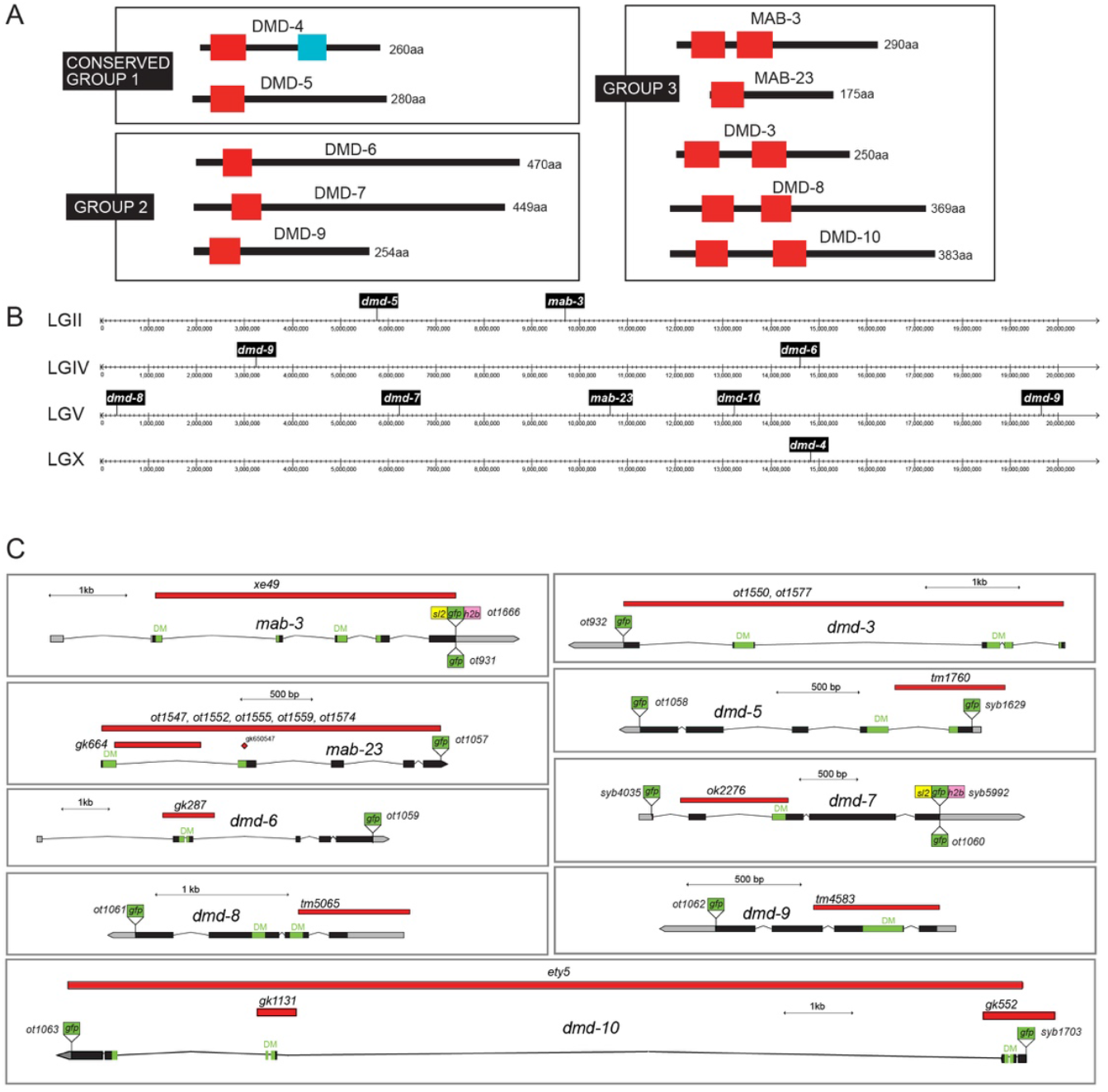
Molecular structures of the *C. elegans* DMRT genes, reporters, and mutant alleles. **(A)** Domain structure. Color codes bear no meaning (from www.wormbase.org). Grouping is based on phylogeny shown in **Fig. S1** and ^12^. *dmd-10* and *dmd-11* were previously annotated as separate genes but, according to conservation and transcript evidence, are now considered a single gene. (**B**) chromosomal location. (**C**) Loci, alleles, and reporters. *dmd-4* CRISPR-tagged allele has been published^21^.

To investigate the evolutionary plasticity of *C. elegans* DMRT genes within the nematode phylum, we examined the DMRT complement in other nematodes, focusing on nematodes with particularly well-sequenced and annotated genomes. In a *Caenorhabditis* species with a different mode of sexual production than hermaphroditic *C. elegans*, the male/female (“gonochoristic”) *C. remanei* species, the complement of DMRT proteins is very similar to that of *C. elegans*, with orthologs of each member of the three *C. elegans* groups shown in **Fig. 1**. One of the orthologues, *dmd-9*, duplicated into two very similar, adjacent paralogs (**Fig. S1, Table S1**).

The hermaphroditic diplogastrid *Pristionchus pacificus*, a more distant relative of *Caenorhabditis* within the clade V subphylum, diverged from *Caenorhabditis elegans* more than 100 million years ago. We still find orthologs of all *Caenorhabditis* DMRT group members in *P. pacificus* (**Fig. S1, Table S1**). However, while there is a single DMD-4 and DMD-5 ortholog, there is a notable expansion of paralogs within the other two groups of DMRT proteins, with a doubling of members of the DMD-6/7/9 group and almost a triplication of the two DM-domain containing group (14 members compared to 5 in *C. elegans*), resulting in a total of 19 *P. pacificus* DMRT genes (**Fig. S1, Table S1**).

In a representative of a different, more basal nematode clade, the clade IV human parasite *Onchocerca volvulus*, a gonochoristic species, we observe the opposite pattern, namely, a smaller set of DMRT proteins compared to *Caenorhabditis* (**Table S1**). One deeply conserved DMRT gene, *dmd-4*, appears to be missing; group 2 is only represented by a single member; but the group 3 DMRT genes with two DM domains appear stable. In the presently incompletely sequenced genome of *Plectus sambesii*, a parthenogenic representative of the even more basal nematode family Plectidae, we found again around 10 DMRT genes (including the conserved *dmd-4* and *dmd-5*), as well as divergent single-DM proteins and several two-DM domain proteins.

Altogether, this analysis indicates that the nematode phylum shows an enlarged, yet flexible, complement of DMRT proteins that is not predicted by its sexual mode of reproduction, including the invention of a two-DM domain configuration apparently entirely restricted to nematodes.

### A complete expression atlas for all *C. elegans* DMRT proteins in both sexes throughout animal development

We set out to systematically characterize the expression patterns of all ten *C. elegans* DMRT proteins.

Because DMRT proteins are known to be regulated both transcriptionally and post-translationally through protein degradation ^17,18,21^, our analysis focused on direct visualization of protein expression through the use of CRISPR/Cas9-engineered reporter alleles of endogenous DMRT gene loci. We previously reported the animal-wide expression of GFP-tagged DMD-4 protein in both sexes, revealing sexually dimorphic expression in the nervous system as well as non-dimorphic expression in other cell types ^21^. Subsequently, in studies investigating the timing mechanisms underlying sexual differentiation, we described the expression of endogenous MAB-3 and DMD-3 reporter alleles, but only in parts of the nervous system ^37^. More recently, the expression of an endogenous DMD-9 reporter allele was characterized in a separate study^38^, and additional reports have examined DMD-3 reporter allele expression with a focus on tail hypodermal cells, as well as DMD-3 and MAB-3 in somatic gonadal cells ^22,23,26^.

For the remaining DMRT genes, previous studies either employed reporter constructs that did not capture the full extent of expression (*mab-23, dmd-5, dmd-6, dmd-10*) or did not examine them at all (*dmd-7, dmd-8*). A recent transcriptomic study indicated sexually dimorphic expression of selected DMRT genes, but lacked independent validation ^39^. Here, using CRISPR/Cas9-generated fluorescent reporter alleles, we present complete expression profiles for all DMRT proteins in both sexes across all developmental stages, including detailed identification of individual neuron classes. Reporter alleles were generated by tagging the endogenous loci at either the N- or C-terminus, or both (**Fig. 1C**). Complete expression patterns are presented in **Table 1, Figures 2-6**, and **Figures S2-S11** and are discussed in the ensuing sections.

**Table 1:** Comprehensive list of DMRT-expressing cells across developmental stages. Parentheses indicate non-neuronal expression. *: DMD-4 expression was previously published ^21^.

|  | PROTEIN | SEX | EMBRYO | TEMPORAL DYNAMICS | ADULT EXPRESSION |
| --- | --- | --- | --- | --- | --- |
| SEXUAL DIMORPHISMS | MAB-3 | male | none | Expression in most neurons starts in late L3/early L4 larval stages; PVW visible from L2; AWA from early L3; transient CEPso glia from late L3/early L4; phasmid socket glia from L3/L4 through adulthood.<br>(Linker cell and pharyngeal muscle visible from late L2/early L3; transient expression in seam cells in L3; neuroblasts and somatic gonad, pharyngeal-intestinal valve cells from L3; hypodermal cells throughout body, intestine, muscle, excretory cell, somatic gonad in L4) | 44 sex-shared neuron classes (143 neurons), 11 male-specific neuron classes (32 neurons) (Fig. 3); head & tail glia.<br>(Tail hypodermis; likely tail muscle; posterior body wall muscle; intestine throughout the body; subset of pharyngeal muscle; pharyngeal-intestinal valve cells; excretory cell; somatic gonad.) |
|  |  | herm | none | (Transient epidermal expression: seam cells and hypodermal cells throughout the body from L3/L4 molt to L4, including P cell descendants.) | None. |
|  | MAB-23 | male | none | Neuronal expression starts in larval stages; PVM expression starts in L3 through adulthood; additional male-specific neurons from L4.<br>(Tail blast and muscle cells from L1; body wall muscle along the body from L2 and stronger in L3; additional neuroblast from L3) | 2 sex-shared neuron classes (PHC, PVM); 6 male-specific neuron classes (Fig. 3).<br>(Body wall muscle.) |
|  |  | herm | none | Transient expression occurs in PHC in L2 & L3; PVM from L3 through adulthood.<br>(very dim in tail blast cells in L1-L2; dim body wall muscle cells in L2, dimmer in L3; two non-neuronal cells above the vulva in late L4) | 1 sex-shared neuron class (PVM). |
|  | DMD-3 | male | none | Neuronal expression starts in late L3/early L4. (Expression in linker cell and other gonadal cells from early L3; neuroblasts, hypodermal cells throughout the body in L3/L4, stronger than hermaphrodites; tail tip hypodermis in L3-L4; body wall muscle visible along the body from L3, stronger in L4.) | 4 sex-shared neuron classes; 15 male-specific neuron classes (Fig. 3).<br>(Body wall muscle along the body; a few somatic gonad cells.) |
|  |  | herm | none | (Expression in anchor cell in L3 and early L4; transient dim expression in hypodermal cells throughout the body in late L3/early L4) | None. |
|  | DMD-4* | male | onset in bean stage | 10 sex-shared neuron classes.<br>(Pharynx; hmc.) | Same as in larvae (Fig. 3), except 2 phasmid neuron classes downregulated.<br>(Pharynx; hmc.) |
|  |  | herm | onset in bean stage | 10 sex-shared neuron classes.<br>(Pharynx; hmc.) | Same as in larvae (Fig. 3).<br>(Pharynx; hmc.) |
|  | DMD-9 | male | onset in 2-cell stage | Mostly 8 sex-shared head neuron classes, occasionally variable with 6-9 classes.<br>(Very dim Z2 & Z3 in L1; dim germline in L2-L4; transient dim tail hypodermis in L4) | Mostly 8 sex-shared head neuron classes (Fig. 3).<br>(Very dim germline.) |
|  |  | herm | onset in 2-cell stage | Mostly 8 sex-shared head neuron classes, occasionally variable with 6-9 classes.<br>(Very dim Z2 & Z3 in L1; transient expression in uterine cells in L4, likely toroid cells; dim germline in L2-L4.) | Mostly 8 sex-shared head neuron classes (Fig. 3).<br>(Likely spermatheca-uterine valve cells; very dim germline.) |
|  | DMD-10 | male | onset in 3-fold stage | Variable expression in AWA, AFD, and AIN; additional neurons visible sporadically (ADL, ASI, ASH, ASJ).<br>(No non-neuronal expression; highly enriched in dauers in all somatic cells.) | Variable expression in AWA, AFD, and AIN; 2 phasmid neuron classes (PHA, PHB); 3 male-specific tail neuron classes (PCC, R4A, dim R6A).<br>(No non-neuronal expression.) |
|  |  | herm | onset in 3-fold stage | Variable expression in AWA, AFD, and AIN; additional neurons visible sporadically (ADL, ASI, ASH, ASJ).<br>(No non-neuronal expression; highly enriched in dauers in all somatic cells.) | Variable expression in AWA, AFD, and AIN; expression in phasmids (PHA, PHB) is either not visible or dimmer than in males.<br>(No non-neuronal expression.) |
| NO SEXUAL DIMORPHISMS | DMD-5 | both | onset in early gastrulation | 2 sex-shared neuron classes (RIM, ASJ) in all larval stages; dim ADL occasionally visible. | 2 sex-shared neuron classes (RIM, ASJ); dim ADL occasionally visible. |
|  | DMD-6 | both | onset around 8-cell stage | Ubiquitous in all somatic cells. | Ubiquitous in all somatic cells. |
|  | DMD-7 |  | onset in 2-cell stage | Expressed in 42 sex-shared neuron classes with some variability in expression levels.<br>(Expressed in all other somatic tissue types.) | Dimly expressed in 42 sex-shared and 11 male-specific neuron classes (Supp Fig. S8).<br>(Expressed in all other somatic tissue types.) |
|  | DMD-8 | both | onset in 3-fold stage | 3 sex-shared head neuron classes (AFD; occasionally dim ASK and RIA).<br>(Pharyngeal muscle and head hypodermis from L1; seam cells from L4.) | 3 sex-shared head neuron classes (AFD; occasionally dim ASK and RIA).<br>(Head hypodermis; seam cells; occasionally pharyngeal muscle.) |

**Figure 2:**
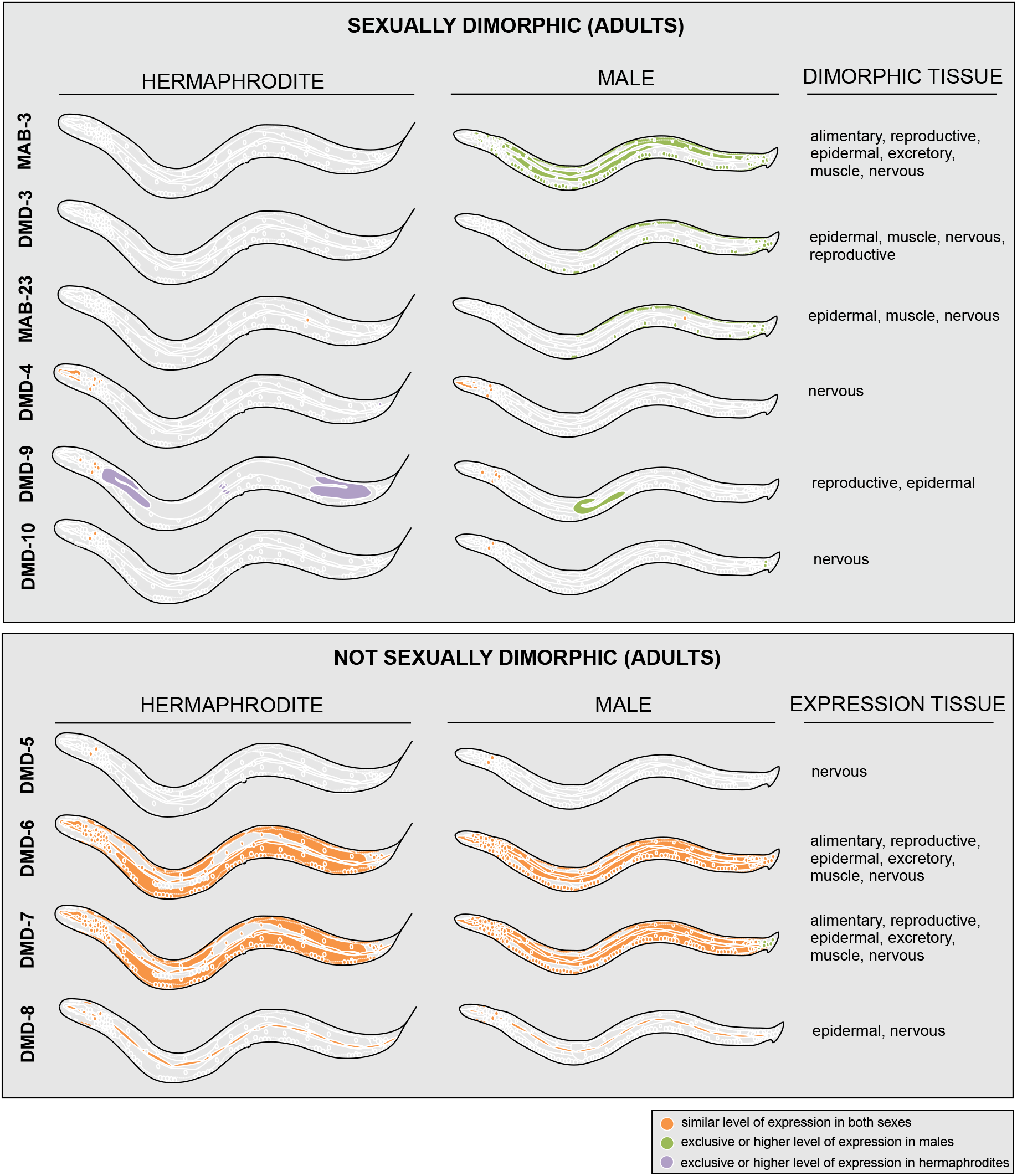
Summary schematics for all DMRT expression patterns in adults. Expression for DMRT reporter alleles is schematized by tissue type. See **Fig. 5, 6** and **Table 1** for details. DMD-7 is considered somatically ubiquitous because its expression is very broad and displays dim fluorescent that is not discernable in some cells (**Fig. S7, S8**).

### Six *C. elegans* DMRT proteins reveal sexual dimorphism in all tissue types

Of the ten DMRT proteins, six exhibited sexually dimorphic expression in adult animals (**Fig. 2-4, Table 1**). In most cases, dimorphisms reflect expression that is either exclusive or enriched in males compared to hermaphrodites, with two exceptions: DMD-4, which shows enriched expression in the phasmid neurons PHA and PHB in hermaphrodites ^21^, and DMD-9, which is expressed in the hermaphrodite somatic gonad (**Fig. 2, 4D**).

Notably, we observe sexually dimorphic DMRT protein expression in all major cell types: neurons, glia, hypodermis, intestine, muscle cells, reproductive organs, and the excretory system (**Fig. 2-4, Table 1**). In the ensuing sections, we describe these patterns, as well as their developmental dynamics.

### DMRT proteins reveal extensive sexual dimorphisms in sex-shared neurons

294 neurons are generated in both *C. elegans* sexes (“sex-shared” neurons)^40–42^, falling into 116 distinct classes. Using the landmark neuron-identification tool NeuroPAL ^43,44^, we systematically characterized the expression of DMRT proteins in all neurons. Of these 294 neurons, 163 express DMRT proteins in a cell-specific manner (excluding the two DMRT genes that are likely ubiquitously expressed, DMD-6 and DMD-7), and 143 (~49% of all sex-shared neurons) exhibit expression of at least one DMRT protein exclusively in one sex (predominantly the male) (**Fig. 3**). This corresponds to sexual dimorphisms in 44 out of 116 sex-shared neuron classes, spanning all three major neuron types—sensory, motor, and interneurons. Male-specific MAB-3 expression alone accounts for much of this dimorphism (**Fig. 3**). In addition to MAB-3, three additional DMRT proteins, DMD-3, MAB-23, and DMD-10, also exhibit male-enriched or male-specific expression in sex-shared neurons (**Fig. 2, 3**). Specifically, DMD-3 is expressed exclusively in the male in motor/interneurons PVN and PDB, the sensory neuron PHC, and two of the eleven AS motor neurons (AS10, AS11) in the ventral nerve cord. MAB-23 shares male-enriched expression with DMD-3 (and MAB-3) in PHC, a morphologically and functionally highly dimorphic sex-shared neuron ^36,45,46^. Co-expression of these specific DMRT proteins is also quite commonly observed in male-specific neurons (see section below). DMD-10 shows male-enriched expression in phasmid sensory neurons PHA and PHB, similar to MAB-3 and opposite to DMD-4, which is expressed in these two neuron classes in hermaphrodites but not males ^21^. Notably, we did not notice any expression of the DMD-10 reporter allele in the AVG neuron, in which we had previously observed male-enriched expression using a 1 kB promoter fusion of the initially mis-predicted gene (*“dmd-11”*)^47^.

**Figure 3.**
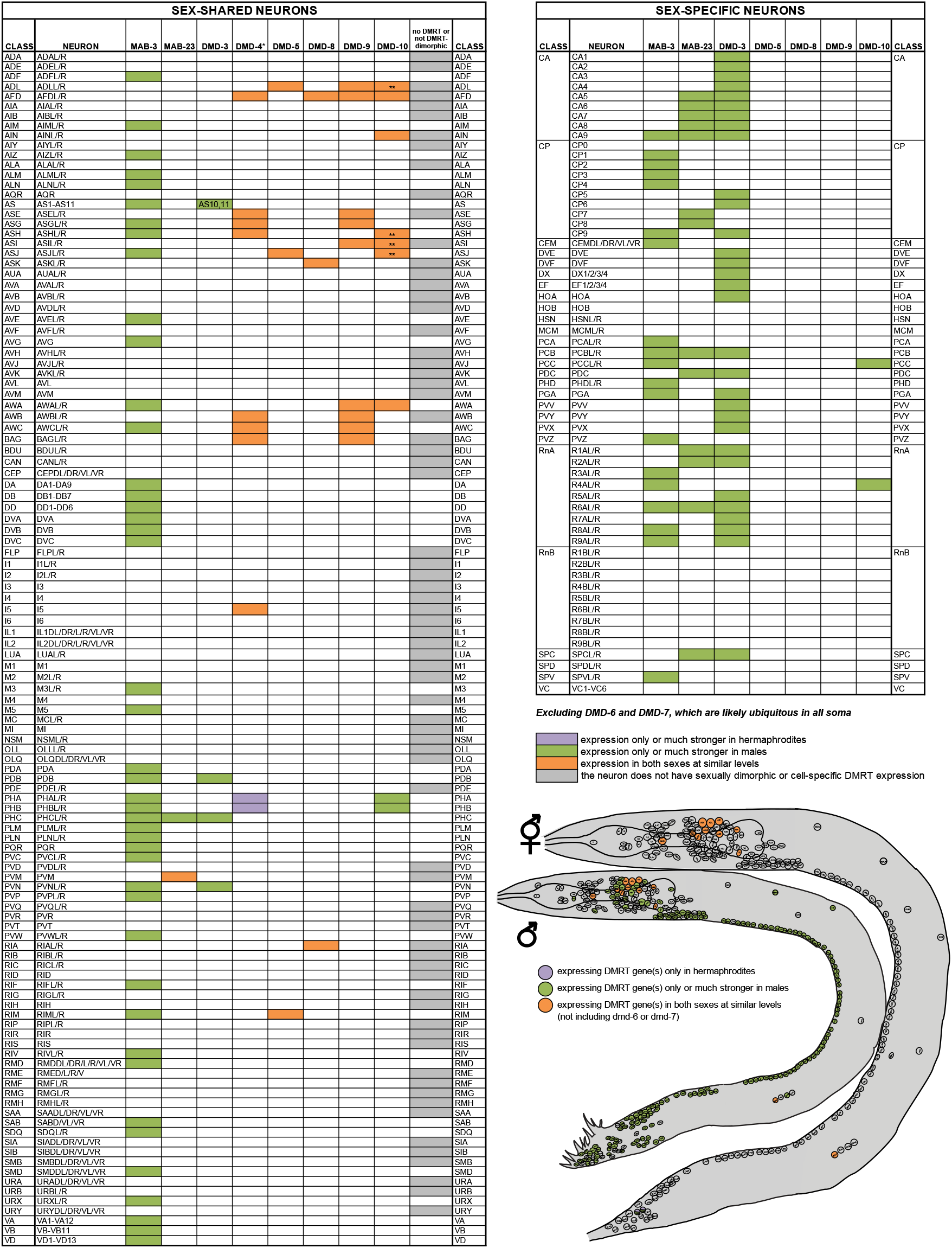
Expression of DMRT proteins in individual neuron classes. Individual neuron classes were identified using the landmark strain NeuroPAL or additional reporters (see Materials and Methods). Tables list sex-shared and sex-specific neurons separately. Worm schematics label neurons according to their anatomical positions. Green: expression exclusive to, or stronger in, males; Purple: expression is exclusive to, or stronger in, hermaphrodites; Orange: expression present in both sexes at similar levels. Gray box in the table for sex-shared neurons indicate neurons that do not express any DMRT proteins (excluding DMD-6 or DMD-7) or whose expression is not sexually dimorphic. DMD-6 is excluded because it is somatically ubiquitous (thus panneuronal). DMD-7 expression is dim in most neurons and is also excluded. Because of its low expression levels and its presence in many neurons, it is possible that DMD-7 could be expressed in even more neurons, maybe even panneuronally, with expression levels too faint to be detected by our microscopes. *DMD-4 expression was previously published ^21^. **Expression visible in larvae.

### DMRT proteins are expressed in most, but not all, sex-specific neurons

In addition to the sex-shared neurons, *C. elegans* also possesses 8 neurons exclusively in hermaphrodites (2 classes, HSN and VC) and 93 in males (“male-specific” neurons). Our analysis revealed no DMRT protein expression in the 8 hermaphrodite-specific neurons, but expression in 69 out of the 93 (74%) male-specific neurons (**Fig. 3, 4**). This encompasses 21 of the 25 male-specific neuron classes (HOB, MCM, SPD, and RnB show no cell-specific DMRT expression). Among these 69 neurons, 22 express more than one DMRT protein, most commonly the combination of DMD-3 and MAB-23 (**Fig. 3**).

**Figure 4.**
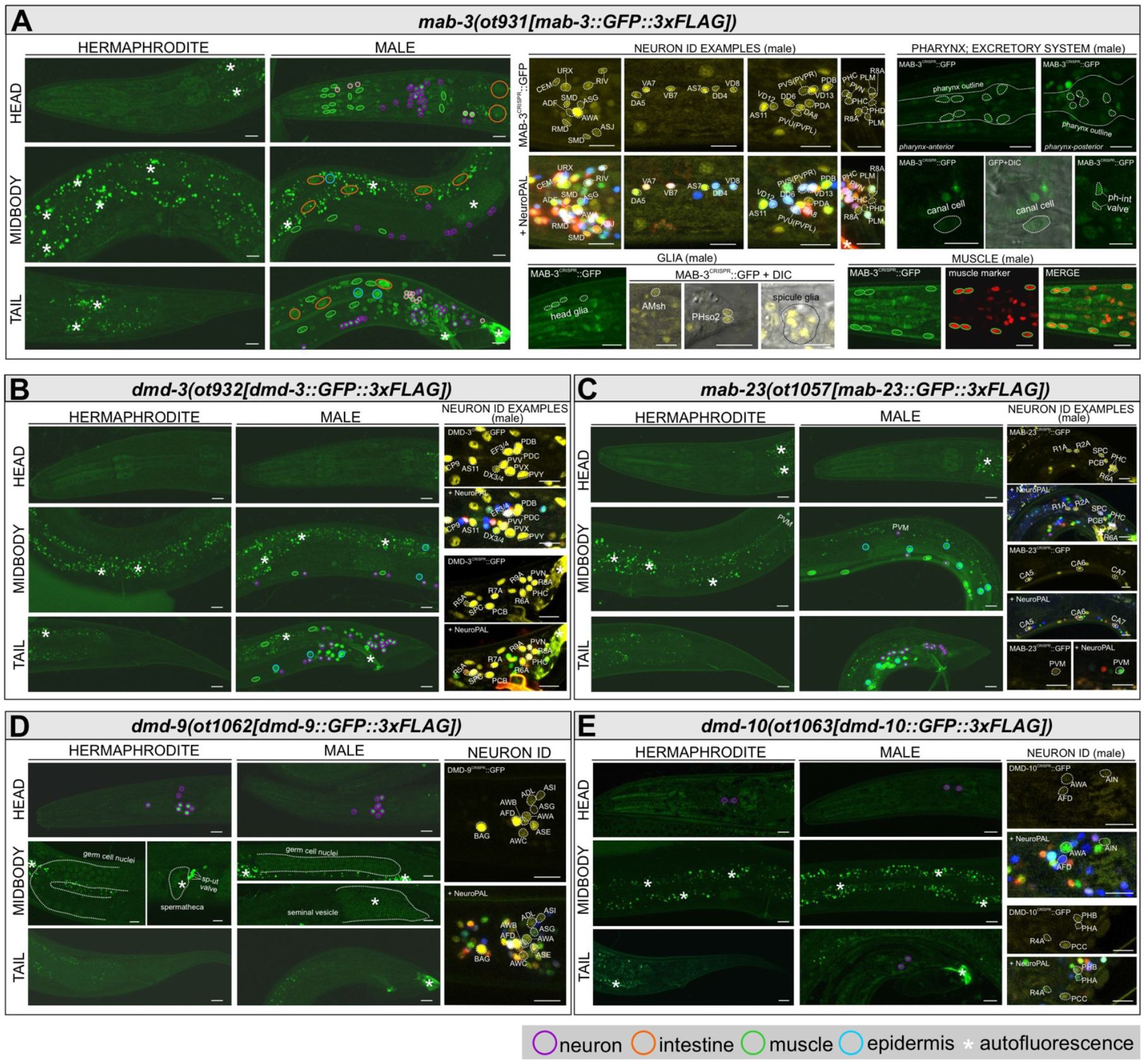
Widespread sexually dimorphic expression of DMRT proteins in adult animals across multiple tissues. Reporter alleles for MAB-3, DMD-3, MAB-23, DMD-9, and DMD-10 exhibit sexually dimorphic expression in adults, with dimorphism largely driven by male-enriched expression. GFP from the reporter alleles shows a yellow pseudo-color in NeuroPAL ID images. (**A-E**) Tissue types are outlined in different colors as indicated in the legend on the figure. For a complete list of neurons, see **Figure 3**. For each gene, overall expression is shown in the left panels (head, midbody, and tail) for both sexes. Right panels show examples of identified neuron classes and other cells in males. (**A**) *mab-3(ot931)* is expressed broadly in males throughout the head, midbody, and tail in a large number of neurons, the intestine, muscle cells, pharyngeal cells, pharyngeal-intestinal valve cells, and the excretory canal cell. No expression is detectable in adult hermaphrodites. Thin, solid line in the top right panels outline the shape of the pharynx. ph-int valve, pharyngeal-intestinal valve cells. (**B**) *dmd-3(ot932)* is expressed in neurons, body wall muscles, and epidermal (hypodermal) cells in the midbody and tail of male animals and is not visible in adult hermaphrodites. (**C**) *mab-23(ot1057)* is expressed in body wall muscles, hypodermal cells, and neurons in the midbody and tail of the male. It is also expressed in neuron class PVM in adult hermaphrodite, in a non-sexually dimorphic manner. (**D**) *dmd-9(ot1062)* is expressed in 8 pairs of head neuron classes (occasionally 6-9 pairs) in both sexes, with neuron classes labeled in the right panels. In the midbody, both sexes show visible, albeit very dim, expression in germ cell nuclei. Additional fluorescent signals sometimes appear in the hermaphrodite spermatheca and male seminal vesicle, but because these regions are prone to autofluorescence, we cannot exclude the possibility that both or one of the signals are in fact autofluorescence (see also **Fig. S10F**). Notably, clear signal is present in cells located between the spermatheca and uterus. The morphology of these cells aligns with the spermatheca-uterine (sp-ut) valve cells (www.wormatlas.org). (**E**) *dmd-10(ot1063)* is expressed in 3 pairs of head neurons in both sexes and in additional neurons in the male tail. Because expression is dim in some neurons, the left panels do not capture all neuronal signals; see the zoomed-in right panels for neuron class identities. Scale bars, 10 μm.

The expression signatures of DMRT proteins in male-specific neurons, in combination with our recently published expression atlases for neurotransmitter pathway genes, neuropeptides, and homeobox genes ^48,49^, help refine anatomically defined neuron classes into molecular distinctive subclasses. This is evident in the CA, CP, and ray neuron classes, each comprising multiple members. For instance, among the 9 CA neurons, CA5-8 co-express MAB-23 and DMD-3, CA1-4 express only DMD-3, and CA9 expresses MAB-3, DMD-3, and MAB-23. This expression pattern is consistent and reinforces previous predictions of CA1-4 representing one subclass and CA9 a distinct subclass ^48,49^. Similarly, within the 10 CP neurons, CP0 is the only one that lacks DMRT protein expression, distinguishing it as a separate subclass. CP1-4 express MAB-3, alongside neurotransmitter genes *unc-17/VAChT* and *unc-47/VGAT* ^48^ and neuropeptide *flp-20* ^49^, defining a unique CP subclass. CP5 and CP6 express DMD-3 and the neurotransmitter genes *eat-4/VGLUT* and *unc-47/VGAT*. CP7 and CP8 express MAB-23, *unc-17/VAChT, unc-47/VGAT*, and neuropeptides *flp-5* and *flp-20*, forming another distinct subclass. Finally, CP9 expresses MAB-3 and MAB-23, neuropeptides *flp-7* and *flp-32*, and homeobox gene *vab-3* ^49^, marking it as a separate subclass.

### A DMRT protein reveals sexual dimorphisms in glia

The DMRT protein MAB-3 also shows sexually dimorphic expression in glia. From as early as the L3/L4 molt, MAB-3 is observed transiently in the sex-shared head glial cells CEPso in male animals (**Fig. 6E**). In the same time window, MAB-3 starts to be expressed in the tail phasmid socket cells PHso1 (**Fig. 6E**). In adult males (and not hermaphrodites), MAB-3 is expressed in sex-shared AMsh, PHso2, and additional head glial cells, as well as in the male-specific spicule glial cells (**Fig. 4A**).

### Sexually dimorphic expression of DMRT proteins in cell types outside the nervous system

We also found sexually dimorphic DMRT protein expression in sex-shared cells outside the nervous system. Male-specific expression of MAB-3 in the intestine has been previously reported and shown to be required for regulating sex-specific yolk protein expression in the intestine^16,33^. We further found that MAB-23 and DMD-3 are expressed dimorphically in body wall muscle cells along the length of male animals (**Fig. 4B, C, S3, S4, Table 1)**. Additionally, transient DMD-3 expression in hypodermal cells along the body during the L3/L4 molt also appears enriched in males (**Fig. S3**). To our knowledge, other than the global sex regulator TRA-1 and the BTB-Zn finger protein EOR-1 described later in this study, no other genes have yet been reported to display sexually dimorphic expression in these sex-shared tissue types.

MAB-3 displays sexually dimorphic expression in three additional cell types not previously known to exhibit morphological or molecular sexual dimorphisms, namely pharyngeal muscle cells, pharyngeal-intestinal valve cells and the canal cell of the excretory system (**Fig. 4A)**. These findings validate the use of DMRT protein expression as a strategy to uncover previously unrecognized sexually dimorphic features in otherwise sex-shared cell types.

DMD-9 displays dim expression in the adult germline of both sexes, as well as in cells most closely align with the spermatheca-uterine valve cells in hermaphrodites (**Fig. 4D**). In larval stages, it also exhibits additional expression in hermaphrodite uterine cells as well as hypodermal cells in the male tail (**Fig. S10D**).

### Non-sexually dimorphic expression of DMRT proteins

Not every DMRT protein shows a sexually dimorphic expression pattern. Three of them—DMD-5, DMD-6, and DMD-8—are robustly expressed but the respective reporter alleles show no discernable sexual dimorphisms throughout development. DMD-5 is expressed in three head neuron classes (RIM, ASJ, and variably in ADL) in both sexes (**Fig. 5A**). Unlike previous promoter-based reporter transgenes ^47^, no male-specific expression was observed in the AVG neuron. We confirmed this pattern with two different reporter alleles, one in which DMD-5 is tagged at the N-terminus, the other at the C-terminus (**Fig. 1C**).

**Figure 5.**
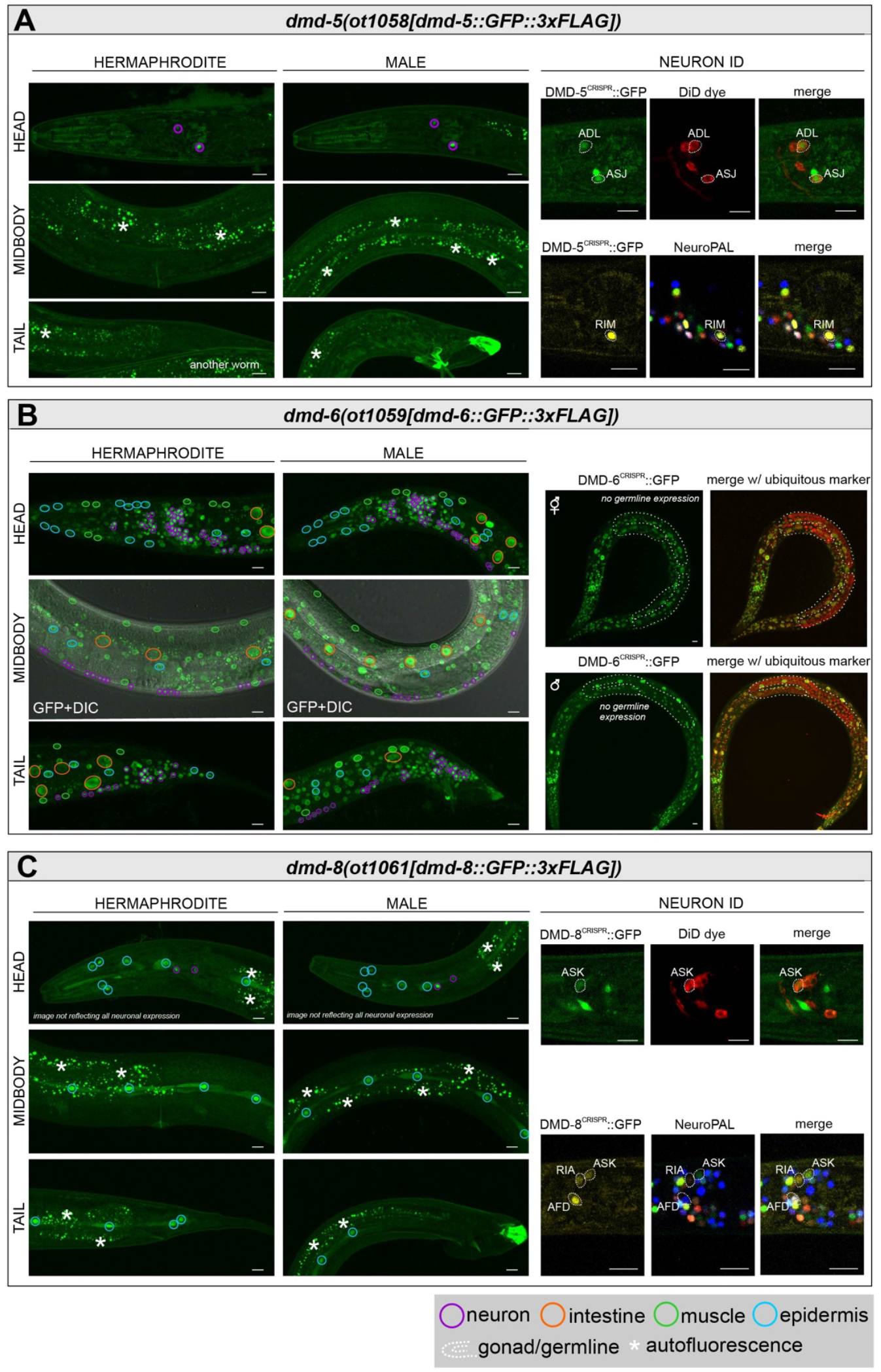
Several DMRT proteins lack sexually dimorphic expression in adults. Reporter alleles for DMD-5, DMD-6, and DMD-8 exhibit non-sexually dimorphic expression in adults. DMD-7 is also considered not dimorphic and is shown in **Fig. S7, S8**. GFP from the reporter alleles shows a yellow pseudo-color in NeuroPAL ID images. (**A-C**) Tissue types are outlined in different colors as indicated in the legend on the figure. For each gene, overall expression in the head, midbody, and tail is shown in the left panels for both sexes. (**A**) *dmd-5(ot1058)* is expressed in 3 pairs of head neuron classes in both sexes (ASJ is not visible in left panel overviews due to low signal). ***(B)*** *dmd-6(ot1059)* is expressed in all somatic cells throughout the body in both sexes. Right panels show an overlay with the ubiquitous marker *mel-28(bq6)*, confirming that expression is absent from in the germline. (**C**) *dmd-8(ot1061)* is expressed in head hypodermal cells and seam cells along the body in both sexes, and in 3 pairs of head neuron classes. Scale bars, 10 mm.

DMD-8 is expressed in a different set of three neuron classes (AFD, and more variably in ASK and RIA), and also in pharyngeal muscle, hypodermal cells, and seam cells along the body (**Fig. 5C**). In striking contrast, DMD-6 is ubiquitous in all somatic nuclei (**Fig. 5B**). Its related paralog, DMD-7, also shows a very broad but much dimmer and, in some cells, hardly discernable expression. Unlike any other DMRT protein, tagged DMD-7 protein also does not appear to be restricted to the nucleus **(Fig. S7)**. Similar patterns of DMD-7 expression and localization are observed with the GFP tag inserted either at the N- or at the C-terminus (**Fig. 1C, S7**). Tagging the 3’ end of the *dmd-7* gene with an *SL2::GFP::H2B* cassette generated brighter signals, corroborating that *dmd-7* is most likely ubiquitously expressed in both sexes **(Fig. S8)**.

Among the DMRT proteins that do show robust, sexually dimorphic expression, MAB-23, DMD-9, DMD-10, along with the previously reported DMD-4 ^21^, also exhibit non-sexually dimorphic expression in other cell types. Specifically, MAB-23 is expressed in the sensory neuron PVM in both sexes from L3 through adulthood (**Fig. 4C, S4**); DMD-9 is typically expressed in eight (occasionally six to nine) neuron classes in both sexes, including neurons not reported earlier (**Fig. 4D**) ^38^; and DMD-10 is expressed in three head neuron classes (albeit variable, **Fig. 4E**).

Together, these non-sexually dimorphic yet distinct expression patterns suggest that DMRT proteins may also play broader roles in animal development that are independent of sexual differentiation.

### Developmental dynamics of DMRT protein expression

DMRT proteins show distinct patterns of onset of expression. The broadly expressed DMD-6 and DMD-7 proteins are visible in very early embryogenesis (**Fig. S6, S8**), followed by DMD-5 expression in early gastrulation (**Fig. S5**), consistent with a previous survey of embryonic transcription factor expression^50^. DMD-4 expression follows at the bean stage, as previously reported^21^. DMD-8 and DMD-10 expression follows at the 3-fold stage (**Fig. S9, S11**). The remaining DMRT proteins, MAB-3, MAB-23, and DMD-3 (all of which highly sexually dimorphic in the adult), show no detectable embryonic expression (**Table 1**).

During larval development, the non-sexually dimorphic DMRT proteins (DMD-5, DMD-6, DMD-8, and the likely non-dimorphic DMD-7) largely maintain their expression in the same cell types across all developmental stages (**Fig. S5-S9**). For DMD-10, which becomes sexually dimorphic in adults, larval expression remains non-dimorphic but occasionally includes additional head neurons in both sexes compared to adults (**Table 1**).

In contrast, MAB-3, MAB-23, and DMD-3, the three DMRT proteins that show the most extensive sexual dimorphisms in adults, display notable developmental dynamics, with dimorphic expression arising progressively across larval stages.

Consistent with previous results based on a transgenic reporter line ^34^, MAB-23 is expressed in epithelial blast cells in the tail at the first larval stage. This expression is stronger in males compared to hermaphrodites (**Fig. S4**).

In the L2 stage, MAB-3 shows expression in the tail neuron PVW in males but not in hermaphrodites (**Fig. 6B, E, Fig. S2**). At the same time, MAB-23 expression in the tail blast cells remains enriched in males (**Fig. S4**). In the nervous system, MAB-23 starts expression in PHC neurons in both sexes (**Fig. S4**), but this expression later diminishes specifically in hermaphrodites, resulting in male-specific maintenance into adulthood. Thus, MAB-3 expression in PVW represents an early-onset neuronal dimorphism, while MAB-23 expression in PHC provides an example of dimorphism arising through the *loss* of expression in hermaphrodites rather than *gain* in males.

**Figure 6:**
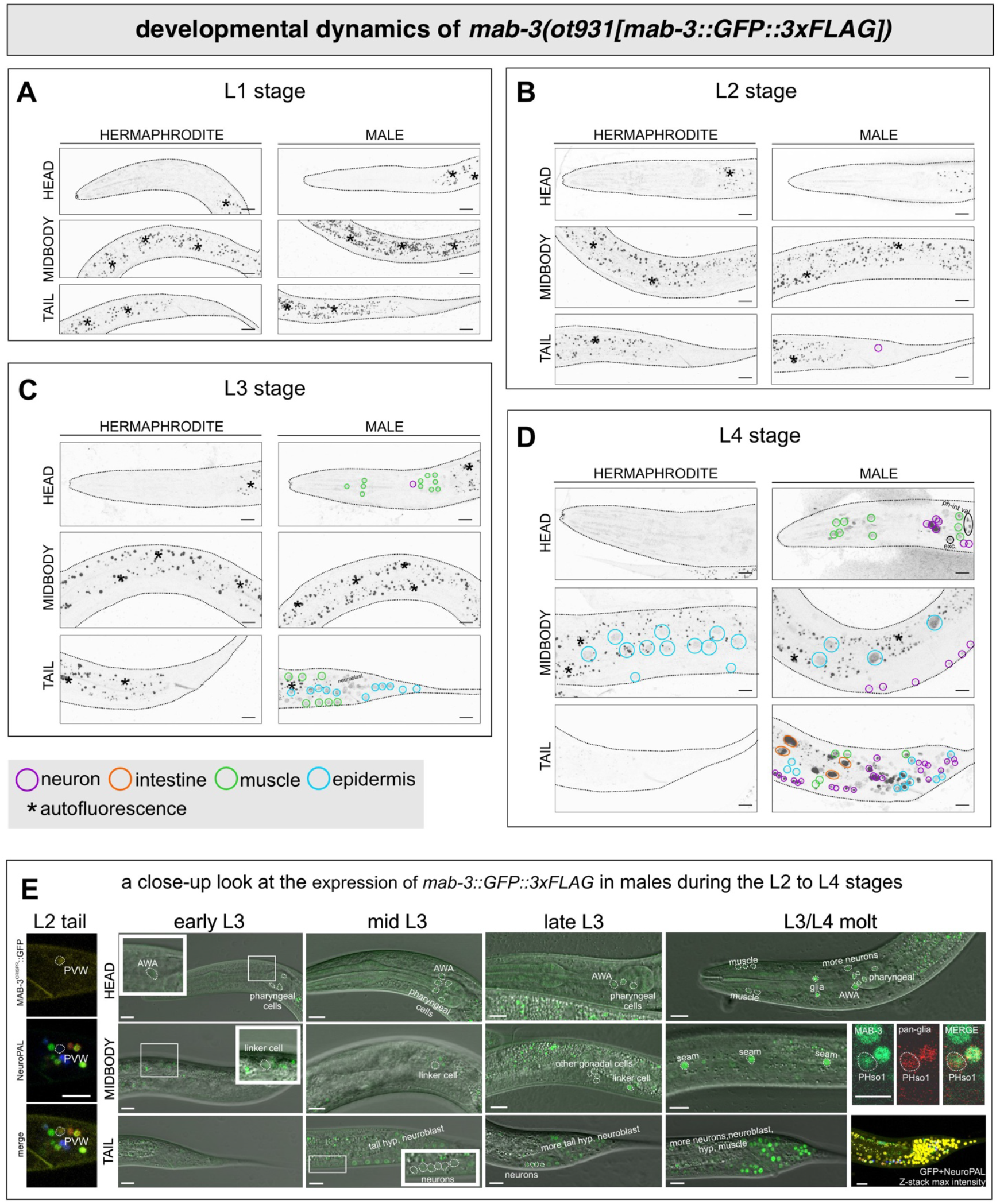
**Temporal dynamics of DMRT expression—the MAB-3 example** *mab-3(ot931)* expression across four larval stages. (**A-D**) overall expression in L1 (**A**), L2 (**B**), L3 (**C**), and L4 (**D**) stages. Tissue types are outlined in different colors as indicated in the legend on the figure. (**E**) Zoomed-in views showing the onset of expression in specific tissue and cell types in males during substages between L2 and L4. Dim expression is shown in insets for selected cells. For glial expression during the L3/L4 molt, a pan-glial marker *otIs870[mir-229p::3xnls::TagRFP]* was used to label glial cells. In the phasmid socket cell images, either of the labeled cells corresponds to PHso1; the other is likely PHso2. To confirm expression in specific cell types, a *mab-3::SL2::GFP::H2B* reporter allele is also analyzed (**Fig. S2**). exc., excretory canal cell; ph-int val, pharyngeal-intestinal valve cells. Scale bars: 5 μm for PHso1 panels; 10 μm for all other panels.

In early L3, another case of early neuronal dimorphism appears: MAB-3 expression in AWA is only detectable in males and not hermaphrodites. Additionally, MAB-3 is visible in pharyngeal muscles in the male (**Fig. 6B, E, Fig. S2**). From early to mid L3, MAB-3 and DMD-3 expression initiates in the male linker cell (**Fig. 6E, Fig. S3**), generally coinciding with the timing reported previously ^22^. Meanwhile, MAB-23 starts to be enriched in males in body wall muscles at this stage (**Fig. S4**). DMD-3 begins expression in additional gonadal cells in the male, as well as in the anchor cell in hermaphrodites (**Fig. S3**). By mid to late L3, DMD-3 also starts to show male-enriched expression in body wall muscle (**Fig. S3**), and MAB-3 shows increasing expression in neuroblasts, additional gonadal cells, and tail hypodermal cells in the male (**Fig. 6C, E**).

From late L3 stage through early L4 (during the L3/L4 molt), four CEPso glial cells start to express MAB-3 exclusively in males (**Fig. 6E**), consistent with a recently reported sexually dimorphic function in these cells^51^. In the same time window, both MAB-3 and DMD-3 continue to be expressed in the linker cell, tail neuroblasts, muscles, and hypodermal cells in the male. Furthermore, MAB-3 is visible in seam cells in both sexes, and DMD-3 shows hypodermal expression along the body in both sexes, with its intensity appearing stronger in males (**Fig. 6E, Fig. S3**). In the tail, DMD-3 displays hypodermal expression in the male tail tip, and its pattern generally aligns with previously reported patterns using an N-terminally GFP-tagged allele for DMD-3 ^23,26^ and earlier reporter transgenes of DMD-3 and MAB-3 ^18^.

At the L4 stage, male-enriched expression of MAB-3, MAB-23, and DMD-3 in neurons reaches its peak (**Fig. 6D, Fig. S3, S4**). Additionally, CEPso glial expression of MAB-3 begins to decline in late L4, representing a rare example of transient and dynamic sexually dimorphic expression. Meanwhile, MAB-3 expression in seam cells gives way to hypodermal cells in both sexes, again with a greater intensity in males (**Fig. 6D**). For MAB-23, expression in PHC diminishes in hermaphrodites during this stage, while it remains robust in males through adulthood (**Fig. S4, Fig. 4C**). In addition, we also observed transient expression of DMD-9 in hypodermal cells in the male tail (**Fig. S10**).

Another notable DMRT protein expression dynamic is observed in the environmentally controlled dauer stage. Among all DMRT proteins, DMD-10 protein expression transforms from a restricted expression pattern in a few cell types (as described above) to a robust expression in all cells of dauer stage animals, starting upon dauer commitment at the L2d stage (**Fig. S11**). This upregulation occurs in both sexes and is reversed upon exit from the dauer stage.

### DMRT genes are broadly required for conferring neurotransmitter identities in male-specific neurons

Having characterized DMRT expression patterns, we next turned to genetic loss of function analyses, focusing on DMRT genes with sexually dimorphic expression in the nervous system. Two *C. elegans* DMRT genes, *dmd-3* and *mab-23*, have previously been shown to regulate cholinergic and dopaminergic identities in type A ray neurons^19,34^. However, it remained unclear whether these effects are restricted to these specific ray neurons or extend to other parts of the male-specific nervous system. Given our new expression data showing that over two thirds of male-specific neurons express at least one DMRT protein, we asked whether DMRT genes generally affect neurotransmitter identities across all male-specific neurons.

A recently completed atlas of neurotransmitter identities in all *C. elegans* neurons for both sexes, generated using CRISPR/Cas9-engineered reporter alleles^48^, enabled us to examine all 93 male-specific neurons for the effects of the three DMRT genes with the most sexually dimorphic neuronal expression, namely *mab-3, dmd-3*, and *mab-23*. Using this resource, in combination with CRISPR/Cas9-engineered null alleles of *mab-3, dmd-3*, and *mab-23* that eliminate all concerns about the null nature of classic, canonical mutant alleles (**Fig. 1C**; see Methods), we found that the three DMRT genes broadly shape neurotransmitter identities in nearly all male ganglia.

In the head, the male-specific CEM sensory neurons are cholinergic^48,52^. Among the DMRT genes, only *mab-3* is expressed in this neuron class. We therefore tested the expression of the acetylcholine vesicular transporter *unc-17/VAChT* in a *mab-3* null mutant. Expression of *unc-17/VAChT* was decreased in *mab-3* null mutant animals (**Fig. 7A**). The CEM neurons have also been reported to express *mod-5*, the serotonin uptake transporter in *C. elegans* ^48^, and its expression was similarly reduced in *mab-3* mutants (**Fig. 7B**). Thus, *mab-3* controls both the cholinergic identity and serotonin uptake capacity of the CEM neurons.

**Figure 7.**
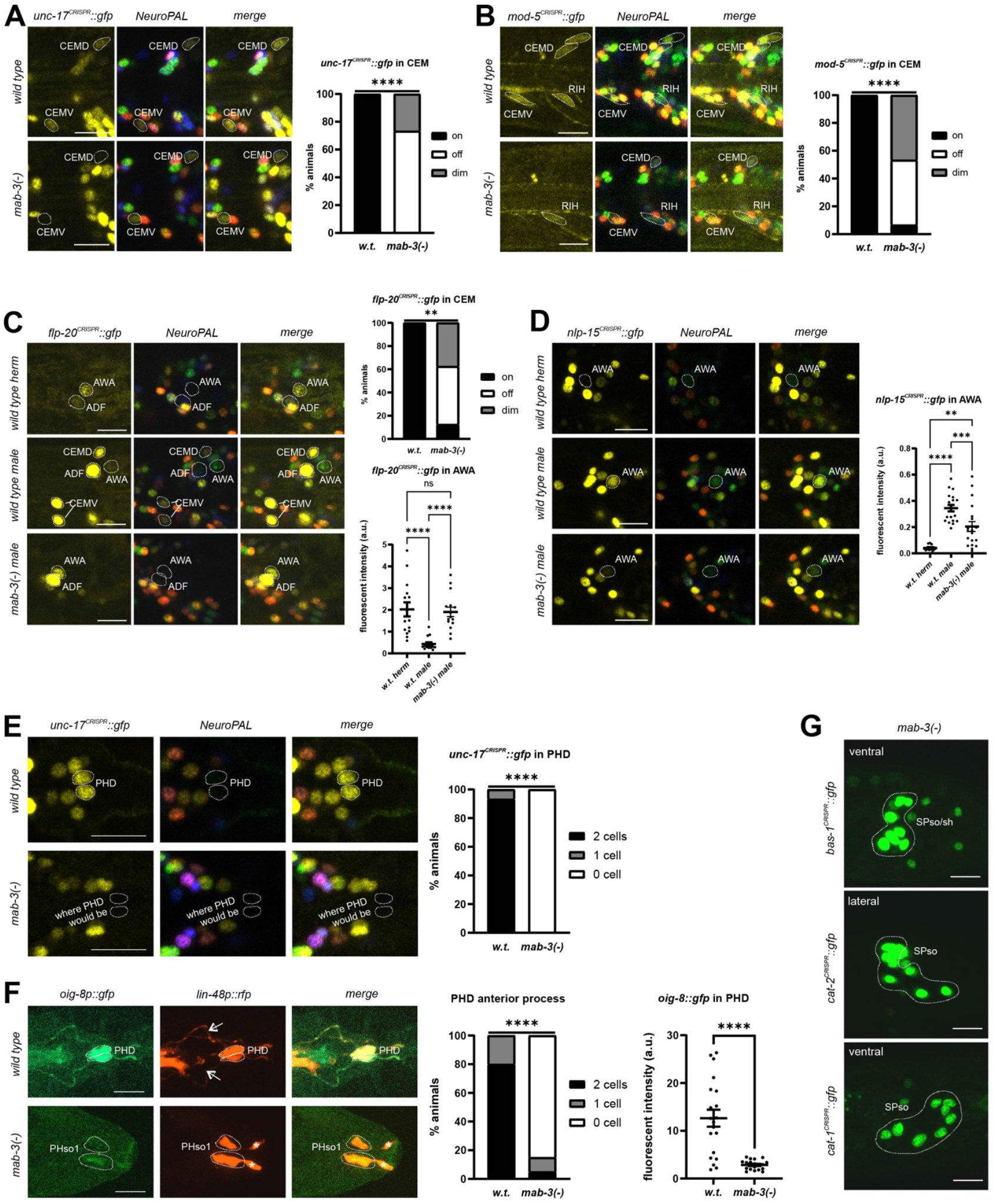
Effects of *mab-3* on neuronal and glial differentiation. The impact of *mab-3(xe49)* null mutant on all main neurotransmitter identities (glutamatergic, cholinergic, GABAergic, and monoaminergic) was examined across all 93 male-specific neurons. Its effects on neuronal differentiation were also analyzed for two neuropeptide genes, as well as markers of glia-to-neuron differentiation and additional glial identities. See also **Table S2**. (**A, B, E**) For neurotransmitter identities, *mab-3(xe49)* downregulates cholinergic identity (**A**, *unc-17(syb4491)*) and reduces expression of the serotonin uptake transporter (**B**, *mod-5(vlc47)*) in head male-specific neuron class CEM. It also downregulates cholinergic identity (**E**, *unc-17(syb4491)*) in the tail neuron class PHD. (**C, D**) For neuropeptide identities, *mab-3(xe49)* downregulates expression of <u>F</u>MRF-like <u>p</u>eptide *flp-20* (**C**, *flp-20(syb4049)*) in the male-specific CEM neurons. Notably, in the sex-shared neuron class AWA, *mab-3* governs a neuropeptide signature switch. In wild type, *flp-20* expression in AWA is enriched in hermaphrodites. *mab-3(xe49)* feminizes male AWA by ***upregulating*** *flp-20* to levels similar to wild type hermaphrodites (**C**). Conversely, expression of *nlp-15* (*nlp-15(syb7428)*) is enriched in males in wild type; *mab-3(xe49)* feminizes AWA by ***downregulating*** *nlp-15*, shifting expression closer to (although not identical to) hermaphrodite levels (**D**). (**F**) *mab-3(xe49)* prevents the PHso1-to-PHD glia-to-neuron transdifferentiation event in males. PHD is labeled by *oig-8* in green (*drpIs4[oig-8p::GFP+pha-1(+)]*) and *lin-48* in red (*drpIs3[lin-48p::tdTomato]*). Arrows indicate PHD neuron-specific anterior axon-like processes; asterisks indicate PHso1 glia-specific sockets. (**G**) *mab-3(xe49)* does not detectably affect effects on the dopaminergic identities (*bas-1(syb5923), cat-2(syb8255), cat-1(syb6486)*) in male spicule glial cells SPso and/or SPsh. Scale bars, 10 μm. Scoring and statistics: see Materials and Methods for scoring criteria. Specifically, fluorescence scoring in CEM neurons used the on/off/dim system; AWA expression was quantified using fluorescence-intensity measurements. Neuron used for normalization: AIZ for *flp-20* and AVA *for nlp-15* expression. For the PHD neuron class, the number of neurons was counted. \*\**P≤0*.*01*, \*\*\*\**P≤0*.*0001*, ns = not significant (A, B, C for CEM, E, F for PHD anterior process: Fisher’s exact test; C for AWA and D: Kruskal-Wallis test followed by Dunn’s multiple comparison test; F for *oig-8* in PHD: Mann-Whitney test).

In the ventral nerve cord, among the nine male-specific CA neurons, all of them are cholinergic, and CA1-CA4 and CA7 are also glutamatergic. Among the ten male-specific CP neurons, CP1-CP4, CP7, and CP8 are cholinergic; CP0, CP5, and CP6 are glutamatergic, CP9 is GABAergic, and CP1-CP6 are also serotonergic^48^. Eight out of the nine CA neurons exhibited altered cholinergic and/or glutamatergic identities upon DMRT gene loss (**Fig. 8A-D, A’-D’**). In *dmd-3* null mutant animals, CA1-CA4 lost expression of *eat-4/VGLUT*, the glutamate vesicular transporter (**Fig. 8A, A’**); in *mab-23* null mutants, CA5 and CA6 gain *eat-4/VGLUT* expression (**Fig. 8B, B’**). In CA7, *dmd-3* null mutant animals lost *eat-4/VGLUT* expression but gained *unc-17/VAChT* (**Fig. 8C, C’**), consistent with a switch from glutamatergic to cholinergic identity. In *mab-23* null mutants, CA7 also gained cholinergic identity. In CA8, *mab-23* and *dmd-3* mutants had opposite effects: *dmd-3* null mutants increased *unc-17/VAChT* expression whereas *mab-23* null mutants decreased it (**Fig. 8D, D’**). Notably, in CA5-CA9, the NeuroPAL landmark strain also showed color shifts, losing its original color and gaining an increased level of the color green (**Fig. 8B-D, B’-D’, Table S2**). These changes suggest that in these neurons, *mab-23* and *dmd-3* likely affect additional genes (harbored in the NeuroPAL transgenes) beyond *unc-17/VAChT* and *eat-4/VGLUT*. Despite DMRT expression in CP neurons, we did not detect changes in their neurotransmitter identities upon DMRT gene loss.

**Figure 8.**
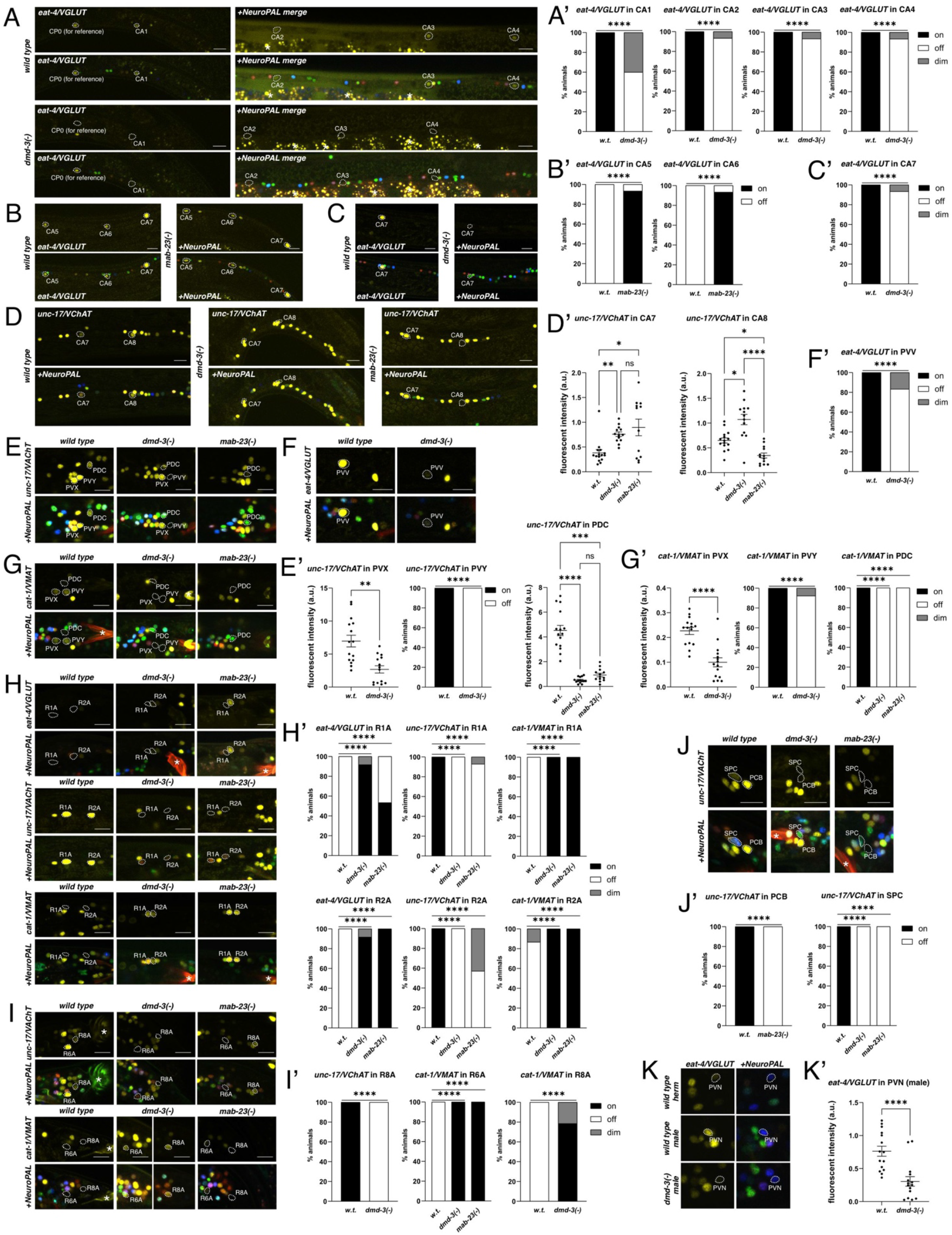
Effects of *dmd-3* and *mab-23* on neuronal differentiation. The impact of *dmd-3* and *mab-23* loss was examined across all 93 male-specific neurons for all main neurotransmitter identities (glutamatergic, cholinergic, GABAergic, and monoaminergic). See also **Table S2**. Mutant alleles were generated by either crossing *dmrt* null mutants or by CRISPR/Cas9-genome engineering when stable homozygous lines could not be obtained by genetic crosses (see Materials and Methods). For glutamatergic identities, *dmd-3(ot1550)* and *mab-23(1552)* were analyzed in *eat-4(syb4257)*; for cholinergic identities, *dmd-3(1577)* and *mab-23(1547)* were analyzed in *unc-17(syb4491)*; for GABAergic identities, *dmd-3(ot1577)* and *mab-23(ot1555)* were analyzed in *unc-25(ot1372)*; for monoaminergic identities, *dmd-3(1577)* and *mab-23(ot1559)* were analyzed in *cat-1(syb6486)*. (**A-D, A’-D’**) In the ventral nerve cord, *dmd-3(-)* downregulates glutamatergic identities in CA1-CA4 (**A, A’**); *mab-23(-)* upregulates glutamatergic identities in CA5-6 (**B, B’**). (**C, D, C’, D’**) In CA7, both mutants upregulate cholinergic identity, with *dmd-3(-)* additionally downregulating glutamatergic identity. In CA8, *dmd-3(-)* upregulates and *mab-23(-)* downregulates cholinergic identity. (**E-G, E’-G’**) In the pre-anal ganglion, cholinergic identities of PDC, PVX, and PVY are downregulated in *dmd-3(-)*, and that of PDC is also downregulated in *mab-23(-)* (**E, E’**). The same pattern is observed for monoaminergic identities (**G, G’**). Additionally, glutamatergic identity of PVV is downregulated in *dmd-3(-)* (**F, F’**). (**H, I, H’, I’**) In the ray neurons, both mutants show the same effects in R1A and R2A neurons by upregulating their glutamatergic and monoaminergic identities and downregulates their cholinergic identity (**H, H’**). (**I, I’**) In R6A, both mutants downregulate cholinergic and upregulates monoaminergic identities, and *dmd-3(-)* also has similar effects in R8A. (**J, J’**) In the cloacal ganglion, both mutants downregulate cholinergic identities in SPC, and *mab-23(-)* also produces a similar effect in PCB. (**K, K’**) In the sex-shared neuron class PVN, the glutamatergic identity is enriched in males in wild type and is decreased/feminized in *dmd-3(-)*. Scale bars, 10 μm. Asterisks in microscopic images represent autofluorescence. Scoring and statistics: see Materials and Methods for scoring criteria. Neurons used for normalization in fluorescence intensity measurements were: DA6 for *unc-17* expression in CA7, VA11 for *unc-17* expression in CA8, HOB for *unc-17* expression in PDC and PVX, HOA for *cat-1* expression in PVX, and the NeuroPAL blue channel for *eat-4* expression in PVN. \**P≤0*.*05*, \*\**P≤0*.*01*, \*\*\**P≤0*.*001*, \*\*\*\**P≤0*.*0001*, ns = not significant (E, G’ for PVX, K’: Mann-Whitney test; D’, E’ for PDC: Kruskal-Wallis test followed by Dunn’s multiple comparison test; all others: Fisher’s exact test).

In the pre-anal ganglion, *dmd-3* affected both cholinergic and monoaminergic identities in three interneurons, PDC, PVY, and PVX. In *dmd-3* null mutant animals, all three showed reduced *unc-17/VAChT* and *cat-1/VMAT* expression (**Fig. 8E, G, E’, G’**). This is notable because these neurons, long thought not to possess monoaminergic identity, were recently characterized as betainergic^48^. The decreased expression of *cat-1/VMAT*, the vesicular transporter for monoamines, suggests that *dmd-3* may regulate this betainergic identity. Similarly, in the PDC interneuron, *mab-23* also affect *unc-17/VAChT* and *cat-1/VMAT* expression, implying that multiple DMRT genes control cholinergic and monoaminergic features in this neuron. Additionally, the glutamatergic identity of PVV was lost in *dmd-3* null mutant animals (**Fig. 8F, F’**). NeuroPAL also showed color changes in PDC and PVV, indicative of a cellular differentiation defects: in both *mab-23* null mutants and *dmd-3* null mutants, PDC gained a green color, while *dmd-3* PVV lost its red color (**Fig. 8E-G, E’-G’, Table S2**). These observations further indicate that *dmd-3* and *mab-23* influence additional genes beyond neurotransmitter markers.

In the lumbar ganglion, previous studies reported that *dmd-3* and *mab-23* affect cholinergic and dopaminergic identities in type A ray neurons^19,34^. Our analyses of *unc-17/VAChT* and *cat-1/VMAT* expression largely confirm these findings, with some specific differences. In both *mab-23* and *dmd-3* null mutants, expression of the *unc-17/VAChT* reporter allele decreased in R1A, R2A, and, R6A, while *cat-1/VMAT* expression increased (**Fig. 8H, I, H’, I’, Table S2**), consistent with loss of cholinergic and gain of monoaminergic (dopaminergic) identity ^19,34^. Additionally, we found that R1A and R2A gained *eat-4/VGLUT* expression in both mutants, indicating that *mab-23* and *dmd-3* regulate the glutamatergic identity of these neurons as well (**Fig. 8H, H’, Table S2**). In R8A, we found a complete loss of *unc-17/VAChT* and gain of *cat-1/VMAT* in *dmd-3* null mutant animals (**Fig. 8I, I’, Table S2**), indicating a cholinergic-to-monoaminergic switch. This extends prior reports that showed a partial increase in *dat-1/DAT* expression ^19^. In contrast to earlier studies reporting identity changes in R3A and R4A ^19^, we did not observe alterations in *unc-17/VAChT* or *cat-1/VMAT* reporter alleles or the expression of the *mab-23* or *dmd-3* reporter alleles in these neurons (**Fig. 3; Table S2**), likely reflecting differences in reporter reagents and neuron identification methods.

The only non-ray neuron class in the lumbar ganglion, PHD, also exhibited altered neurotransmitter identity in a DMRT mutant. In *mab-3* null mutant animals, PHD lost their cholinergic identity (**Fig. 7E**). This neuron is unique in that it is derived from transdifferentiation of the phasmid socket glia PHso1 ^53^. We describe how this process is regulated by *mab-3* in a later section.

Finally, in the cloacal ganglion, the PCB neurons lost their cholinergic identity in *mab-23* null mutants, while SPC neurons lost its cholinergic identity in both *mab-23* and *dmd-3* null mutant animals. In SPC, the NeuroPAL landmark color was also lost and replaced with a green color in both mutant backgrounds (**Fig. 8J, J’; Table S2**).

Among the 69 male-specific neurons expressing *mab-3, dmd-3*, and/or *mab-23*, 30 exhibit regulation of cholinergic, glutamatergic, and/or monoaminergic identities by the DMRT genes, spanning 10 out of the 21 DMRT-expressing neuron classes. We did not observe altered GABAergic identity in the male-specific neurons. In none of the DMRT-regulated cases does the respective DMRT mutant affect the overall generation of the respective neurons. This is notable in light of the expression of MAB-23 in blast cells that generate many of the male-specific neurons; this expression appears to be inconsequential for generation of neurons from these blast cells. To what extent individual DMRT proteins may coordinately control terminal identity features of a neuron beyond its neurotransmitter identity will require further analysis.

### A DMRT gene is required to define neurotransmitter identities in sexually dimorphic, sex-shared neurons

Two vesicular neurotransmitter reporters, *eat-4/VGLUT* and *unc-47/VGAT*, are upregulated in specific sex-shared neurons in the male ^48^. Increased expression of *eat-4/VGLUT* in the PHC neuron class likely relates to a substantial, sex-specific increase in synaptic outputs of PHC in males, and we previously described this so-called “scaling” phenomenon to be dependent on *dmd-3* ^36^. Here, we found a similar process to apparently occur in the PVN interneuron class. These neurons also substantially rewire synaptic connections upon sexual maturation, including an increase in synaptic output (**Fig. S12**)^46^, which we found to correlate with an upregulation of *eat-4/VGLUT* expression in PVN ^48^. Analyzing *eat-4/VGLUT* expression in a *dmd-3* null mutant background, we found that as in PHC, this upregulation depends on *dmd-3* (**Fig. 8K, K’**).

### A DMRT gene is required for a sex-specific neuropeptide signature switch

Beyond sexual dimorphisms in neurotransmitter identities, *C. elegans* also displays pronounced sexual dimorphisms in neuropeptide expression, as revealed by transcriptomic analyses and reporter allele-based expression maps ^39,49,54,55^. One striking example is observed in the sexually dimorphic AWA neuron class ^56^, in which the two neuropeptide-encoding genes *flp-20* and *nlp-15* are expressed in AWA in opposite, sexually dimorphic patterns ^49,54,55^. Using CRISPR/Cas9-engineered reporter alleles for the two neuropeptides, we confirmed that AWA displays a sexually dimorphic split in neuropeptide fate: *flp-20* is enriched in hermaphrodites, whereas *nlp-15* is enriched in males (**Fig. 7C, D**). In *mab-3* null mutants, both identities shift, and in opposite directions: the expression of *flp-20* in males is *gained*, while that of *nlp-15* is *reduced*. Thus, *mab-3* functions as a bidirectional regulator, promoting one neuropeptide identity while repressing another, thereby generating a sex-specific neuropeptide signature switch.

We also found that *flp-20* expression in the male-specific CEM neurons^49,54^ is diminished in *mab-3* null mutant animals (**Fig. 7C**), indicating that *mab-3* also contributes to male-specific neuropeptide identity across multiple neuron classes.

### A DMRT gene is required for sex-specific glia-neuron transdifferentiation

Most sex-specific neurons are generated by cell-specific proliferation of blast cells ^41^. A notable exception is the male-specific neuron class PHD, which arises through direct transdifferentiation from the sex-shared phasmid socket glia, PHso1 ^53^. Because PHD loses its cholinergic identity in *mab-3* null mutant animals, we asked whether this reflects a role of *mab-3* in only specifying neurotransmitter identity in PHD, or it may instead be indicative of *mab-3* controlling the entire PHso1-to-PHD transdifferentiation process.

A hallmark of a successful transdifferentiation is the emergence of a PHD axon that projects anteriorly toward the pre-anal ganglion in the male animal, accompanied by retraction of the PHso1 socket process, which is replaced with a short, dendrite-like posterior projection^53^. To assess whether this transformation occurs in *mab-3* mutant males, we used reporters for *oig-8*, which is expressed in PHD, and *lin-48/OVO1*, which is expressed in both PHD and PHso1^53,57,58^, and examined both morphology and reporter expression. Strikingly, *mab-3(-)* males fail to elaborate the characteristic anterior PHD axon and instead retain socket-like structures typical of PHso1 (**Fig. 7F**). Moreover, *oig-8p::gfp* expression is strongly reduced, mirroring the reduction observed for the *unc-17* cholinergic reporter (**Fig. 7E, F**). Thus, *mab-3* is essential for the PHso1 glia to undergo transdifferentiation, rather than simply modulating PHD neuronal identities.

Because MAB-3 is also expressed in male-specific glia, including spicule socket (SPso) and sheath (SPsh) glia (**Fig. 4A**), we asked whether it also regulates their identities. These cells are known to express dopaminergic pathway genes *cat-1/VMAT, cat-2/TH*, and *bas-1/AAAD* in all or subsets of them ^48,59–61^. However, in *mab-3* null mutant males, expression of these dopaminergic markers appears unaffected (**Fig. 7G**).

### DMRT genes and male-mating behaviors

Male mating defects have previously been reported upon loss of each DMRT gene that normally shows robust sexually dimorphic expression (*mab-3, mab-23, dmd-3, dmd-4, dmd-10*) ^16,19,21,34,47^. However, since we found that the *dmd-10* reporter allele showed a different expression pattern than a *dmd-10* promoter fusion (then called *dmd-11* ^47^), and since that previous work also used an allele that we now understand to only eliminate a part of the gene ^47^, we re-assessed *dmd-10* function. To this end, we generated an unambiguous null mutant in which the entire locus is deleted through CRISPR/Cas9 genome engineering (**Fig. 1C**). *C. elegans* male mating consists of a stereotyped sequence of steps: responding upon contact with a hermaphrodite, moving backwards the body, turning around, locating the vulva, inserting spicules, and ultimately transferring sperm ^62^. We tested male mating behavior and found a non-significant reduction in the male’s ability to locate the vulva, with a frequency similar to that previously reported (**Fig. S13A**)^47^. Neurons previously demonstrated to be involved in mate-contact-induced backward locomotion include the PHA and PHB neurons ^47,63^, which show enriched DMD-10 expression in males (**Fig. 4E**). Thus, we also scored *dmd-10* mutant males for their ability to initiate backward movement in response to contact. We found a significant impairment (**Fig. S13B**). This defect can be partially rescued by expressing DMD-10 specifically in the PHB sensory neurons (**Fig. S13B**). We further tested whether two sexually dimorphic synaptic connections involving these neurons are altered in *dmd-10* null mutants. In wild type animals, PHB>AVG synapse number is increased in males, whereas PHA>AVG synapse number is increased in hermaphrodites ^45–47^. Visualizing synaptic connections using either GRASP^64^ or iBlinc^65^, we found that in *dmd-10* null mutant males, neither connection shows a detectable change in sexual dimorphism compared to wild type animals (**Fig. S13C, D**). These findings suggest that *dmd-10* promotes the transition from initial sensory detection to behavioral engagement during male mating, independently of AVG’s sexually dimorphic synaptic connectivity.

### *C. elegans* BTB-Zn finger proteins display sexually dimorphic expression patterns in somatic and reproductive tissues

In *Drosophila*, the function of the DMRT protein Doublesex is closely linked with that of another transcription factor, Fruitless, a member of a distinct family of transcription factors, the BTB-Zn finger family. ^3,8,9^. Three other BTB-Zn finger proteins, Lola, Tramtrack, and Kruppel are also involved in sexual differentiation in arthropods ^3, 68,62,63^. Hence, we sought to examine whether *C. elegans* BTB-Zn finger proteins may also show signs of involvement in sexual differentiation. The *C. elegans* genome encodes two BTB-Zn finger genes. One of these, EOR-1, was very recently reported to exhibit hypodermal expression in the male and identified as a potential cofactor for DMD-3 in male tip morphogenesis ^26^, but whether its expression is sexually dimorphic was not known. Using a fosmid-based reporter transgene, we found broad EOR-1 expression throughout the animal and a clear male-enriched pattern throughout the hypodermis along the body (**Fig S14A, B**).

We tagged the other, previously uncharacterized BTB-Zn finger protein (F10B5.3), which we named *btbz-2*, with GFP, using the CRISPR/Cas9 system. In L4 stage animals, BTBZ-2 is expressed in hermaphrodite somatic gonadal cells and, very dimly, several head neurons in both sexes (**Fig. S14C**). In adults, it continues to express very dimly in head neurons in both sexes and the hermaphrodite somatic gonad (**Fig. S14D**). Taken together, the sexually dimorphic expression of *C. elegans* and fly BTB-Zn finger proteins indicates that these factors may constitute broadly conserved regulators of sexual differentiation.

## DISCUSSION

We present here the first whole-animal view of DMRT protein expression in a developing and a fully differentiated multicellular organism, defining the extent by which DMRT proteins specify sexually dimorphic features of an animal and revealing specific sets of functions during sex-specific neuronal differentiation processes.

### DMRT expression uncover widespread sexual dimorphisms

We found that six of the ten DMRT proteins are expressed in a sexually dimorphic manner. These six proteins span all three groups of DMRT proteins, including both deeply conserved and nematode-specific family members. The cellular resolution offered by *C. elegans*, with its completely mapped nervous system in both sexes, allowed us to distinguish DMRT expression in sex-specific versus sex-shared tissues with single cell resolution. Several DMRT proteins are expressed in cells that are only present in either males or hermaphrodites, including in reproductive organs, as well as in three quarters of male-specific neurons. Some DMRTs display sexual dimorphic expression in sex-shared tissues, most prominently in almost half of all sex-shared neurons, but also in glia, body wall muscle, hypodermis, pharyngeal muscle, the intestine, the pharyngeal-intestinal valve, and the excretory cell.

Our analysis of genome-engineered reporter alleles substantially expanded and refined expression patterns inferred from previous reporter-based approaches, many of which were based on transgenic promoter-reporter fusions that likely lacked relevant *cis*-regulatory elements. For example, the endogenous *mab-3* reporter allele revealed expression in almost half of sex-shared neurons in males but not hermaphrodites, whereas earlier non-endogenous reporters detected expression only in a few head neurons. Conversely, engineered reporter alleles do not corrobrate some previously reported patterns, such as the *mab-23* head neuron expression described by Lints et al. using an extrachromosomal reporter transgene ^34^. Nor did we detect expression of *dmd-3* in a phasmid neuron, as reported with a multicopy reporter transgene ^18^. Likewise, previous promoter-based reporter transgenes indicated male-specific expression of *dmd-5* and *dmd-10* (formerly *dmd-11*) in the AVG interneuron ^47^, which we could not detect at any stage with our *dmd-5* and *dmd-10* reporter alleles.

With the caveat that reporter alleles may produce expression levels below the limits of detectability, our comprehensive expression analysis of all DMRT family members suggests that not all cells located within sexual dimorphic tissues—be they sexual organs or sex-specifically generated neurons—express a DMRT protein (other than two ubiquitously expressed DMRT proteins). Conversely, not all sex-shared cells known to exhibit molecular or synaptic dimorphisms express a cell type-specific DMRT protein.

The global regulator of sexual identity, TRA-1, is strongly or exclusively sex-biased across most somatic tissues ^69^. TRA-1 has been demonstrated to control the dimorphic expression of several DMRT genes (*mab-3, dmd-3*, and *dmd-4*)^17,18,69^. Given their apparently more restricted expression, DMRT genes should therefore be viewed as one of many TRA-1 effectors, rather than the sole conveyors of TRA-1 function. This is also consistent with the notion that while TRA-1 controls the generation of all sex-specific neuron classes, DMRT genes have only limited roles in patterning sex-specific lineages and instead act primarily to control terminal differentiation events.

### DMRT expression outside the nervous system

Our DMRT expression analysis revealed novel molecular sex differences outside the nervous system, some of which providing potential regulatory explanations for previously suggested functional dimorphisms of these cells. This includes the male-specific expression of MAB-3 in pharyngeal muscles. Although hermaphrodites and males exhibit similar pharyngeal pumping rate under normal conditions on food, males reduce pumping during copulation and display sexually dimorphic pumping in response to the neuropeptide *lury-1*, which itself is not dimorphically expressed ^70,71^. MAB-3 is perhaps required to interpret a non-sex-specific LURY-1 signal in a sex-specific manner.

The sexually dimorphic expression of MAB-23 and DMD-3 in body wall muscle is also a novel molecular sexual dimorphism in this cell type. While there are male-specific muscles in the tail associated with the copulatory apparatus ^41^, sexual dimorphisms in body wall muscle along the length of the worm have only been inferred on a functional level, through elegant cell-autonomous sex-reversal experiments ^72^. These experiments revealed that the sexual identity of muscle contributes to body wave speed ^72^. Sexually dimorphic expression of these two DMRT proteins in body wall muscle may be involved in this phenomenon. Additionally, male-specific expression of MAB-3 in various sex-shared motor neurons reported in this study suggests that MAB-3 could also control well-documented aspects of sex-specific locomotory behavior via the nervous system ^72^.

Male-specific expression of MAB-3 in the excretory canal may explain a recently reported sexual dimorphism in resistance to external stress conditions ^73^. The canal cell is required to tolerate changes in osmolarity of the environment ^74–76^ and males have been shown to be more resistant to detrimental changes in the osmotic environment ^73^. Whether male-specific MAB-3 expression in the canal cell may endow animals to modulate its ability to withstand osmotic challenge will require experimental validation.

In reproductive tissues, we observed DMRT expression in the hermaphrodite anchor cell, uterine cells, and in male linker and other gonadal cells. Germline expression was minimal, limited to faint DMD-9 expression in both sexes. This pattern resembles that of *doublesex* (*dsx*) in *Drosophila*, which is expressed primarily in the somatic gonad, nervous system, and other somatic tissues, with little evidence for strong germline expression ^77–80^. In contrast, the mammalian DMRT1 gene functions in both the somatic gonad and male germline (reviewed in ^81^).

### Complex temporal dynamics of DMRT protein expression

The onset of sexually dimorphic DMRT expression both within and outside the nervous system is notably diverse, appearing as early as the L2 stage and extending through the L3 and L4 stages. This is remarkable because at least two DMRT genes, *mab-3* and *dmd-3*, are regulated by the ubiquitously expressed heterochronic RNA-binding protein LIN-41, which binds to the 3’UTR of *mab-3* and *dmd-3* mRNA to control their degradation ^82^. This 3’UTR-mediated regulation occurs downstream of the miRNA *let-7*, which in turn represses LIN-41 ^83^. The level of mature *let-7* rises between L3 and L4 stages ^83^. This timing corresponds well to the onset of most male-enriched DMRT expression, for example in the sex-shared neurons. However, the earlier appearance of DMRT proteins in other cell types, such as selected neurons and muscle, indicates exceptions to this alignment. Such early-onset expression may therefore reflect cell type-specific differences in LIN-41’s ability to regulate DMRT transcripts, or distinct temporal dynamics of LIN-41 and *let-7* activity across different tissue types.

### DMRT genes broadly regulate neuronal communication in the male nervous system

Equipped with recently mapped molecular markers for male-specific neurons, we focused our mutant analyses of DMRT genes on sex-specific neurons, particularly in the male-specific nervous system (93 neurons in 25 cardinal classes). This focus stemmed from the limited understanding of developmental patterning of the male-specific nervous system. While the first studied *C. elegans* DMRT genes (*mab-3, mab-23, dmd-3*) were discovered through their male tail morphogenesis defects, the functions of these genes have thus far been characterized only in subsets of male-specific neurons, most notably ray sensory neurons ^17,18,33,34,59^. Here, using the neuron-identification tool NeuroPAL and CRISPR/Cas9-generated null mutant alleles that have the entire coding sequences of the DMRT genes removed, we substantially expanded the functional analysis of these three DMRT proteins to the very diverse set of male-specific neurons throughout the male tail, as well as other parts of the nervous system.

Using neurotransmitter identity as a key marker for the generation and proper differentiation of male-specific neuron classes, we corroborated the impact of *mab-3, mab-23*, and *dmd-3* on A-type ray sensory neuron differentiation, largely in agreement with previous studies ^19,34^. We discovered differentiation defects in many additional neuron classes of DMRT mutants, across almost all male ganglia, including male-specific head sensory neurons (CEM neurons in *mab-3* mutants), different subclasses of ventral cord neurons (CA neurons in *dmd-3* and *mab-23* mutants), and several different classes of sensory, motor, and interneurons in the tail across distinct ganglia (interneuron PDC and sensory-motor neuron SPC in *mab-23* and *dmd-3* mutants; PVV motor neuron and PVX and PYX interneurons in *dmd-3* mutants; PCB sensory-motor neuron in *mab-23* mutants). NeuroPAL signals confirm that most of the affected neurons are generated, except for the V lineage-derived ray neurons in *mab-3* null mutants, which is known to affect patterning of these ray lineages^16^. Hence, DMRT proteins are generally not required for the generation of sex-specific neurons.

The similarity of phenotypes of *dmd-3* and *mab-23* mutants in many, but not all, neuron classes is consistent with the possibility that these proteins may act as heterodimers, a mechanism described for vertebrate DMRT proteins ^84,85^.

Our finding that individual DMRT genes affect neurotransmitter identity in some neurons but not others demonstrates that the contribution of DMRTs to neuronal differentiation is highly dependent on cellular context. This context-dependency likely reflects combinatorial interactions between DMRT proteins and other neuronal identity regulators—for example, homeodomain transcription factors, which we found to play a role in the differentiation of several male-specific neuron classes^49^.

In sex-shared neurons, we found that the scaling of glutamatergic signaling (upregulation of *eat-4/VGLUT* in males) of the interneuron PVN, necessitated by synaptic rewiring, depends on *dmd-3*, mirroring the scaling function of *dmd-3* in another sex-shared neuron class, PHC^36^. In contrast, in the sensory neuron AWA, another DMRT gene caused a striking neuropeptide switch: in *mab-3* null mutants, a neuropeptide normally enriched in hermaphrodites became upregulated in males, whereas a male-enriched neuropeptide was downregulated. This reciprocal change is reminiscent of the “neurotransmitter switch” in the interneuron AIM, where neurotransmitter identity transitions from glutamatergic to cholinergic during sexual maturation of males, a process dependent on the male-specific transcription factor *lin-29* ^37,52^.

### A DMRT gene regulates glia-to-neuron transdifferentiation in the male nervous system

Our neuronal differentiation analysis revealed that *mab-3* is required for the male-specific glia-to-neuron transdifferentiation of the PHso1 socket glia into the PHD neuron. An analogous role has been reported in mammals: Dmrt5 regulates the neuron-glia fate switch in the mouse hippocampus^86^. Here, this “switch” refers to the sequential transition in vertebrate neural stem cell competence, from generating neurons prior to producing glia, so that neuronal circuits are established before glial networks form (reviewed in ^87^). Although the vertebrate process involves a change in progenitor output rather than direct transdifferentiation of an already differentiated cells (as is the case for PHso1-to-PHD), both systems point to a potentially universal role of DMRT family members in regulating neuron-glia fate decisions across species.

### DMRT functions beyond sexual differentiation

The non-sexually dimorphic expression of several DMRT proteins suggests that these factors may also have roles unrelated to sexual differentiation, including in embryonic patterning. This idea is supported by the embryonic lethality of *C. elegans dmd-5* mutants ^47^ and various mouse DMRT knock-out studies^8,10,11,13^. However, it is important to note that many previous studies on DMRT function only examined one sex. A recent study found that depletion of the non-dimorphically expressed vertebrate DMRT2 uncovers sex-specific phenotypes in the mouse brain ^28^. Translated to *C. elegans*, it will be interesting to see whether, for example, the non-sexually dimorphic expression of MAB-23 in the PVM neuron class may serve a similar role, for instance antagonizing potential sexually dimorphic programs that might otherwise be induced by factors such as TRA-1.

Another intriguing function of a DMRT gene is suggested by the striking induction of DMD-10 in all cells of the animal upon entry into the dauer stage. While we did not observe any obvious defects in dauer formation of *dmd-10* null mutants, it is conceivable that *dmd-10* may have a role in other aspects of specialized dauer physiology.

### Implications for DMRT gene function in other organisms

Our findings in *C. elegans* suggests that a broad, systematic analysis of DMRT genes in other organisms may uncover previously unknown sex-specific cells or sexual dimorphisms in sex-shared cells. Moreover, many distinct tissue types in mammals, including humans, exhibit extensive sexual dimorphisms in their transcriptomes^1^, yet the regulatory mechanisms underlying these differences are poorly understood and may involve DMRT proteins.

The same holds true for the nervous system. In more complex, vertebrate nervous systems, the extent of sexual dimorphisms remains incompletely explored. DMRT genes may provide a powerful entry point for identifying and dissecting sexually dimorphic regulatory mechanisms. Recent attempts have been undertaken in the mouse brain ^29^. However, since in situ mRNA hybridization (or scRNA analysis) does not capture post-transcriptional regulatory events, which are known to control DMRT protein expression in *Drosophila* and *C. elegans* ^17,18,21^, it will be important to extend transcript-based studies to protein-based studies.

### Sexual dimorphisms beyond DMRT proteins

Our organism-wide analysis of DMRT proteins reveals that they do not mark all cells and tissues that exhibit sex-specific features, indicating that other factors regulate sex-specific gene expression in these cells. In DMRT-negative, sexually dimorphic cells, the master regulator of sexual differentiation, TRA-1, may directly regulated sexually dimorphic effector genes. Other sexually dimorphic transcription regulators exist as well. These include the Zn finger transcription factor LIN-29, a TRA-1 target, which is expressed in a male-specific manner in a subset of sex-shared neurons ^37^, as well as the BTB-Zn finger factor EOR-1, which we describe here to display sexually dimorphic expression in the skin. Other than skin reorganization during copulatory organ formation in the male tail ^23^, worm skin cells along the entire body were not previously known to display sexual dimorphisms, while sexual dimorphism has been documented in human skin ^88^.

The recurrent deployment of BTB-Zn finger transcription factors in sexual differentiation in both flies^3^ and worms (this paper) raises interesting questions about BTB-Zn finger proteins in mammals. The BTB-Zn finger family expanded to almost 50 genes in mammals, and the numerous invertebrate precedents for dimorphic BTB-Zn finger expression and function may stimulate investigations into whether these transcription factors regulate sexual dimorphisms in mammals as well ^89^.

## ACKNOWLEDGMENTS

We thank Chi Chen for assistance with *C. elegans* microinjections to generate transgenic strains, Chien-Po Liao for sharing strain OH20048, Esther Serrano-Saiz for discussing unpublished data, Quinten Cremers for helping with *dmd-10* mutant analysis, Julia Wittes from Iva Greenwald’s lab for the *ckb-3* reporter, and members of the Hobert lab for comments on the manuscript prior to its submission. Some strains were provided by the Caenorhabditis Genetics Center (CGC), which is funded by NIH P40 OD010440.

## AUTHOR CONTRIBUTIONS

Conceptualization: C.W. and O.H. Methodology: C.W. and Y.S. Investigation: C.W. and Y.S. Visualization: C.W. Y.S. M.O.-S., and O.H. Supervision: O.H. and M.O.-S. Writing—original draft: C.W. and O.H. Writing—review & editing: C.W. Y.S. M.O.-S., and O.H.

## DATA AND MATERIAL AVAILABILITY

All data are present in the paper and supplementary materials. All new strains are deposited at the CGC.

## MATERIALS AND METHODS

### *C. elegans* strains and maintenance

Worms were maintained on nematode growth media (NGM) plates at 20°C. *Escherichia coli* (OP50) bacteria was used as a food source. Males were obtained by crossing reporter alleles into *him-5(e1490)* for genes not on Chromosome V and *him-8(e1489)* for those that are on Chromosome V. Strains used in this study are listed in **Data S1** and deposited to the Caenorhabditis Genetics Center (CGC).

### CRISPR/Cas9-based genome engineering and molecular cloning

Reporter alleles *mab-23(ot1057), dmd-5(ot1058), dmd-6(ot1059), dmd-7(ot1060), dmd-8(ot1061), dmd-9(ot1062)*, and *dmd-10(ot1063)* were generated by C-terminally tagging the endogenous loci as previously described ^90^. *mab-3(ot931)* and *dmd-3(ot932)* were published previously^37^. The reporter alleles *dmd-5(syb1629), dmd-7(syb4035), dmd-7(syb5992)*, and *dmd-10(syb1703)* were obtained from SunyBiotech.

The *mab-3* null mutant allele *xe49*, a full locus deletion, was previously described ^91^. To generate the *dmd-3(ot1577)* and *mab-23(ot1574)* null mutant alleles, recombinant Cas9 (IDT), tracrRNA (IDT), a *myo-2::tagRFP* injection marker, and two crRNAs flanking either the *dmd-3* or *mab-23* locus were injected into N2 hermaphrodite gonads. The repair templates were ssODNs. TagRFP-expressing F1 progeny were screened, and deletion alleles were first identified by abnormal male tail morphologies, followed by PCR and sequencing to confirm the mutant loci.

To generate the *dmd-10(ety5)* null mutant allele, Cas9 (IDT) was injected into N2 gonads with tracrRNA (IDT) and three crRNAs, one targeting the *dpy-10* locus ^92^ and two flanking the *dmd-10* locus. The repair template was an ssODN. Rol/dpy F1 progeny were screened by PCR to identify the *dmd-10(ety5)* deletion allele.

To generate DMD-10 rescue constructs, *dmd-10* cDNA was amplified from N2 library. Gibson assembly was used to connect *dmd-10* cDNA with a *gpa-6* plasmid backbone to generate PHB-specific rescue constructs.

For sequences for individual sgRNAs and primers for cloning repair templates, see **Data S2**.

### Expression analysis and cell identification

Expression of reporter alleles was first analyzed without any markers, and the alleles were then genetically crossed into marker strains to aid in identifying specific cell types. In cases where reporter expression was dim, fluorescence patterns were compared with those of reporter alleles alone to rule out the possibility that weak “signals” arose from bleed-through of the marker into the GFP channel. When autofluorescence was a concern, for example in the intestine, germline, and tissues near the male spicule, a control *him-5* or *him-8* strain was used for comparison.

For analysis in adults, animals were separated by sex at the L4 stage and imaged 6 to 9 hours later at adulthood. For neuronal identification (ID), reporter alleles were crossed into the NeuroPAL landmark strains (*otIs669* for *mab-3, dmd-5, dmd-6*, and *dmd-9* alleles; *otIs696* for *dmd-3, dmd-7, dmd-8*, and *dmd-10* alleles). Neuron ID followed published protocols ^43,44^. GFP from the reporter alleles shows a yellow pseudo-color in NeuroPAL ID images. Occasionally, dye-filling with DiD (Thermo Fisher Scientific) was used to confirm the ID for amphid neurons. For glial ID, a pan-glial reporter *otIs870[mir-228p::3xNLS::TagRFP]* was used, and the identity of each glial cell was subsequently determined by stereotypical positional features.

For tissue ID outside the nervous system, cells were recognized first based on morphology using Nomarski optics and by their anatomical position. To ID hypodermal expression, reporter alleles were crossed into a hypodermal marker, *stIs10166[dpy-7p::his-24::mCherry+unc-119(+)]*. Additionally, to distinguish body wall muscle from hypodermal expression, a strain that labels both tissue types with distinct fluorescent colors (SD1546) ^93^ was used to train the experimenter to recognize the characteristic positions and morphologies of each tissue type. For sexing dauers, *arTi112[ckb-3p::mCh::h2b]* was used to label the somatic gonad, and worms were sexed based on the sexually different expression of *ckb-3*.

### Neuronal identity mutant analysis

For neurotransmitter identity analysis, DMRT mutant alleles were crossed into reporter alleles for *unc-17/VChAT* (*syb4491*), *eat-4/VLUT* (*syb4257*), *unc-25/GAD* (*ot1372*), *cat-1/VMAT* (*syb6486*) all in the NeuroPAL background. Additionally, *mod-5/SERT* (*vlc47*) was used for analyzing the effects of *mab-3* null mutants in CEM neurons ^48^. In cases where crosses did not yield stable homozygous lines because the genes were located nearby on the same chromosome, the DMRT mutant alleles were recreated by injecting the same CRISPR injection mix used to generate their null alleles directly into the neurotransmitter reporter strains in the NeuroPAL background. These include: *dmd-3(-)*: *ot1550*; *mab-23(-)*: *ot1547, ot1552, ot1555*, and *ot1559*. See **Fig. 8** for details.

For neuropeptide identity analysis, the *mab-3(xe49)* null mutant ^91^ was crossed into reporter alleles for *nlp-15* (*nlp-15(syb7428[nlp-15:SL2:GFP::H2B]*) and *flp-20* (*flp-20(syb4049[flp-20::SL2::GFP::H2B]*) with NeuroPAL in the background.

Male animals were separated from hermaphrodites at the L4 stage and imaged 6 to 9 hours later together with control neurotransmitter or neuropeptide reporter strains. Neuron ID was performed as described above. For analyzing fluorescent expression in mutant male tails, images were captures from multiple angles (lateral left, lateral right, ventral, and dorsal) to ensure accurate neuron ID, since *mab-3, mab-23*, and *dmd-3* mutants exhibit abnormal tail morphologies. 15-30 animals were scored for each strain. When expression was clearly present or absent, mutant expression was qualitatively scored using three categories: on (signal clearly abundant), off (signal clearly absent), and dim (signal present but visibly reduced relative to control animals). When expression levels were not clearly distinguishable, fluorescence intensity was quantified in ZEN software. For normalization, expression in an adjacent neuron unaffected in the respective mutant was used (see also Legends). For expression in the PHD neuron class, the number of PHD neurons was counted (0, 1, or 2). Statistical analyses were performed using GraphPad Prism version 10.

### Microscopy

For expression and mutant analyses, animals were anesthetized with 50-100 mM sodium azide and mounted on thin, 5% agarose pads on glass slides. They were imaged as Z-stacks using a 40x objective on either a Zeiss LSM880 or LSM980 confocal microscope. Images were processed in ZEN to generate orthogonal projections ^94^. In some cases, brightness and contrast were adjusted uniformly across control and experimental groups in ZEN or CorelDraw to ensure dim expression was visible on the figures.

For transsynaptic labeling experiments, animals were imaged using a 63x objective on a Zeiss LSM880 confocal microscope. Puncta were quantified by examining the original full Z-stack for distinct fluorescent dots in the region where the processes of the two neurons overlap.

### Mating behavior assays

Mating assays were performed as described previously ^47^. Early L4 males were transferred to fresh plates and kept apart from hermaphrodites until they reached sexual maturation. Single virgin males were assayed for their mating behavior in the presence of 10– 15 adult *unc-31(e928)* hermaphrodites on a plate covered with a thin fresh *E. coli* OP50 lawn. Mating behavior was scored within a 15-minute time window or until the male ejaculated, whichever occurred first. Mating was recorded using a Zeiss Axiocam ERc 5s mounted on a Zeiss Stemi 508. Movie sequence was analyzed and males were scored for contact response and vulva location efficiency ^95^. Contact response requires tail apposition and initiation of backward locomotion. Percentage response to contact = 100 x (the number of times a male exhibited contact response/the number of times the male makes contact with a hermaphrodite via the rays)^47,62^. Vulva location efficiency ^96^, or the ability to locate the vulva, was calculated as 1 divided by the number of passes or hesitations at the vulva until the male first stops at the vulva or until the end of the recording. All statistical analyses for mating assays were computed using GraphPad Prism version 10.

