## Supplementary Material for "A panoramic view of the expression and function of the Doublesex/DMRT gene family in *C. elegans*"

**This PDF file includes:**

Figs. S1 to S14
Tables S1 to S2

**Other Supplementary Materials for this manuscript include the following:**

Data S1 to S2

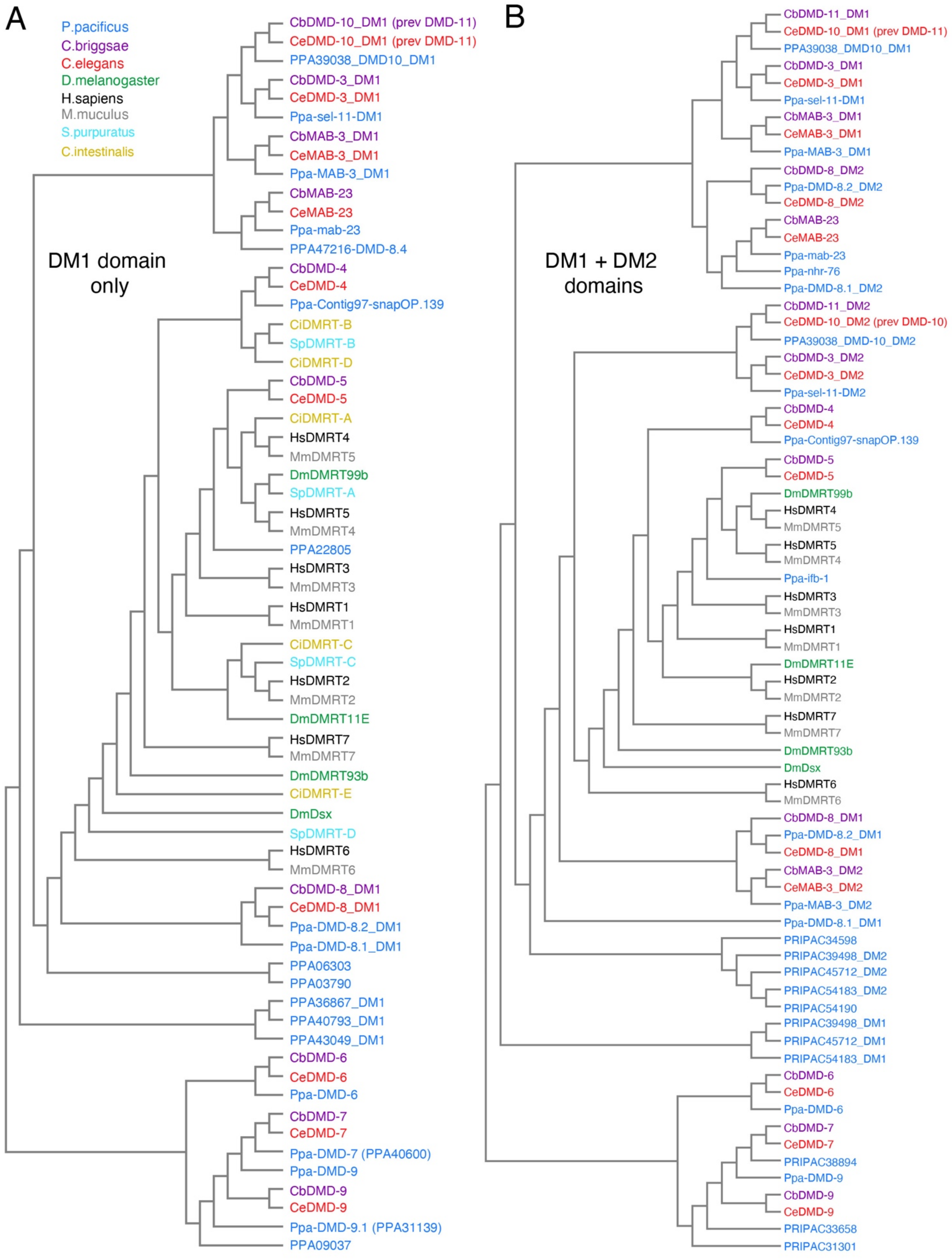

**Figure S1. Phylogeny of DMRT proteins**

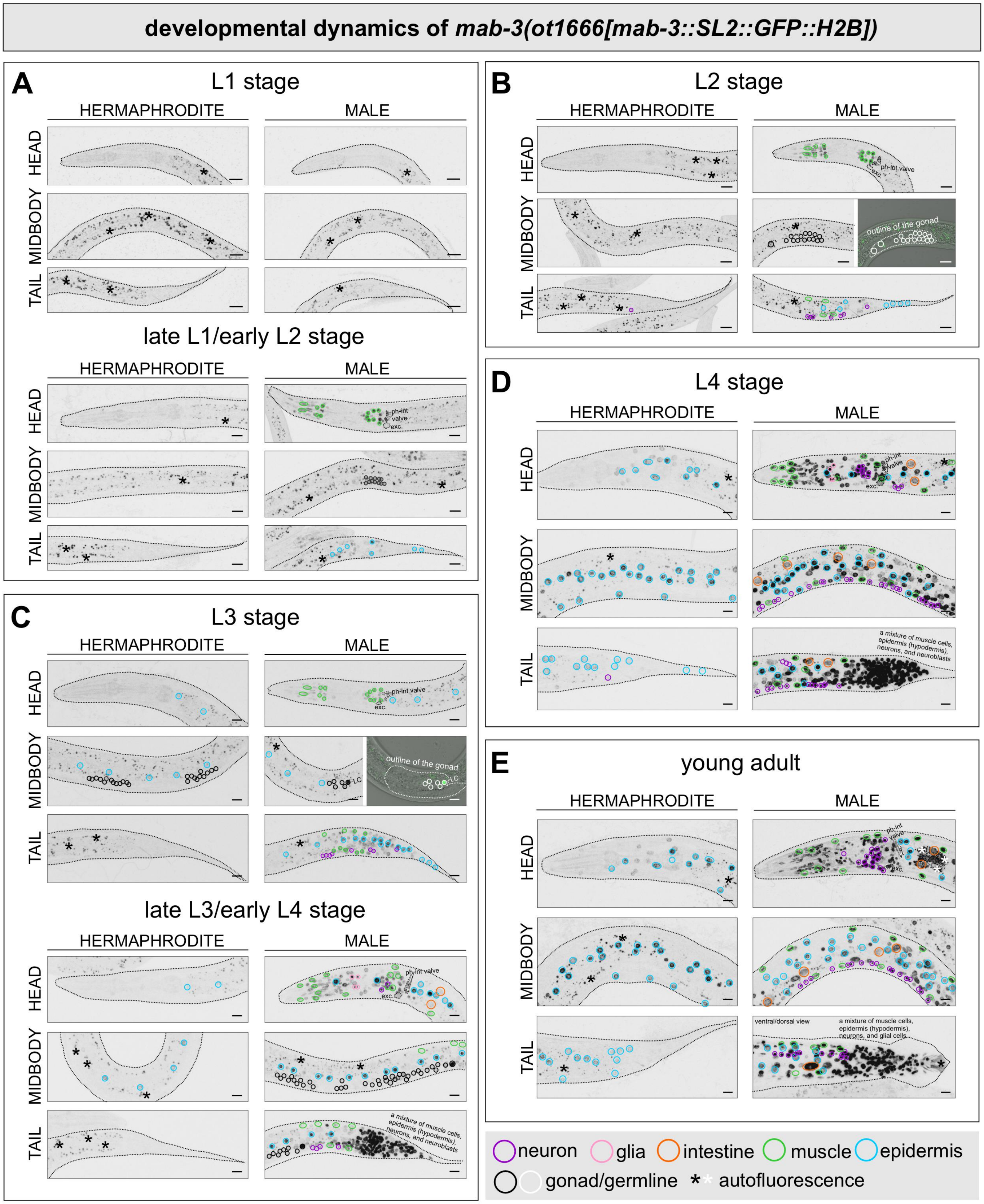

**Figure S2. Expression of an additional *mab-3* reporter allele.** Apart from examining the expression of a reporter allele in which *GFP* is directly fused to *mab-3*, we also examined the expression of an *SL2::H2B::GFP*-based *mab-3* reporter allele, *ot1666* (**Fig. 1C**), across larval and adult stages in grayscale. (**A-E**) overall expression in L1 and late L1/early L2 (**A**), L2 (**B**), L3 and late L3/early

L4 (**C**), L4 (**D**), and young adult (**E**) animals. Tissue types are outlined in different colors as indicated in the legend on the figure. The overall spatial pattern of *mab-3(ot1666)* closely matches the protein expression observed with the *mab-3(ot931)* reporter (**Fig. 4A, Table 1**). As expected from the inclusion of an H2B element, *mab-3(ot1666)* accumulates higher fluorescent intensity, which makes certain features easier to visualize, without altering the sites of expression defined by *ot931*, in which *GFP* is directly fused to *mab-3*. The increased brightness contributes to several practical differences in the visible readout: 1) earlier detection in larval stages, with signal apparent from late L1/early L2 in the pharynx and additional neurons from L2 onward; 2) more readily detectable gonadal and excretory cell expression; 3) visible hypodermal expression in adult hermaphrodites, whereas *ot931* shows hypodermal signal primarily at L4; 4) clearer visualization of a ventral cord neuron in both sexes and more neurons in the male from L2. At this point it is difficult to distinguish whether the earlier onset of the *SL2::H2B::GFP*-based *mab-3* reporter allele, most notably in the excretory cell and pharynx, is simply the result of greater sensitivity of such a reporter cassette or indicative of additional layers of post-transcriptional gene regulation. Dotted lines indicate the outlines of the animals. Cells in the male tail at L3/L4, L4, and adult stages are not individually labeled because they are densely packed and difficult to distinguish accurately. exc., excretory canal cell; ph-int val, pharyngeal-intestinal valve cells. Scale bars, 10 μm.

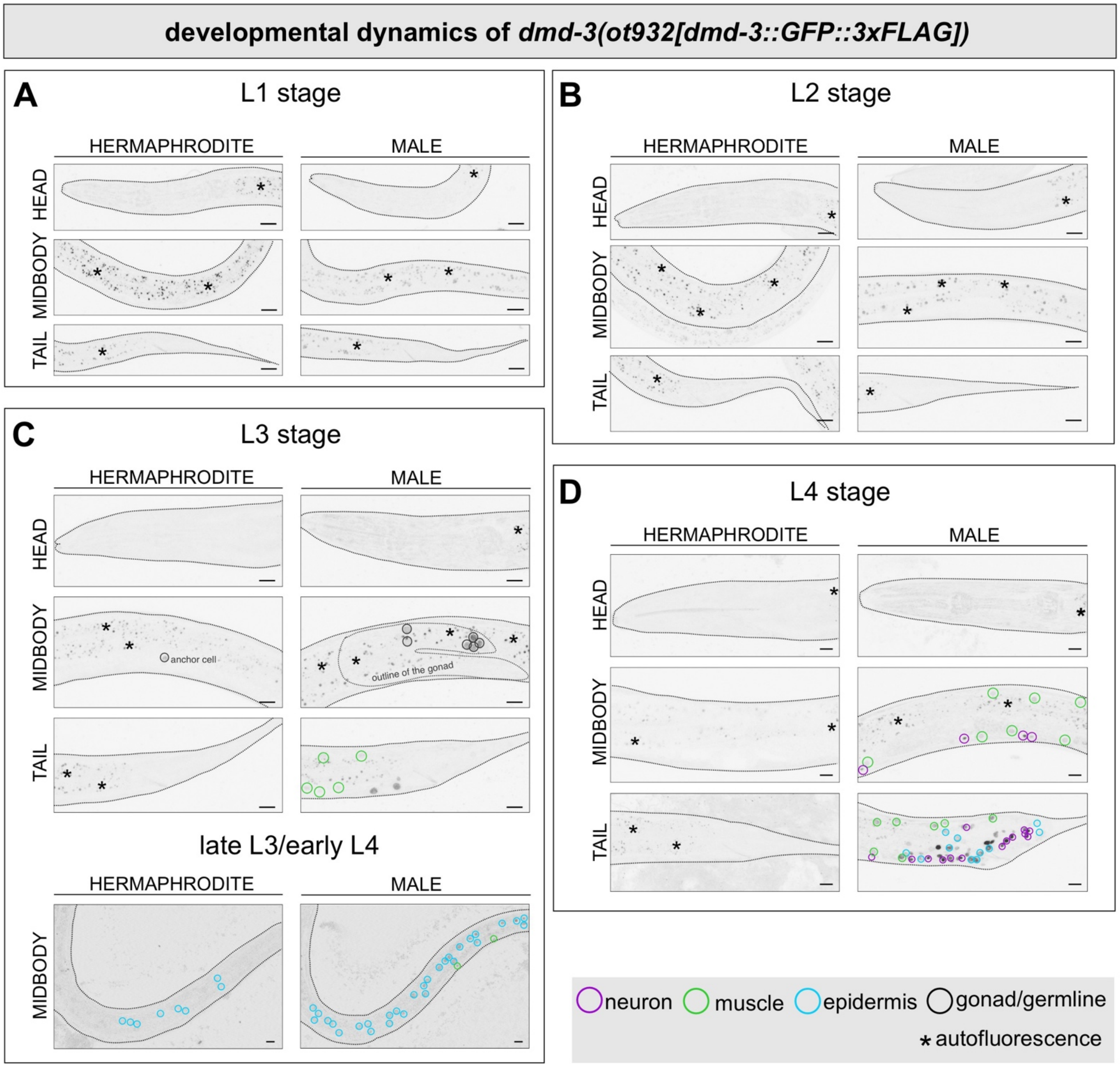

**Figure S3. Temporal dynamics of DMD-3.** *dmd-3(ot932)* reporter allele expression across larval and adult stages in grayscale. (**A-D**) overall expression in L1 (**A**), L2 (**B**), L3 and late L3/early L4 (**C**), and L4 (**D**) stages. Tissue types are outlined in different colors as indicated in the legend on the figure. Refer to **Table 1** and Results for description of expression patterns. Dotted lines indicate the outlines of the animals. Dotted line inside the L3 male animal indicates the shape of the gonad. Scale bars, 10 µm.

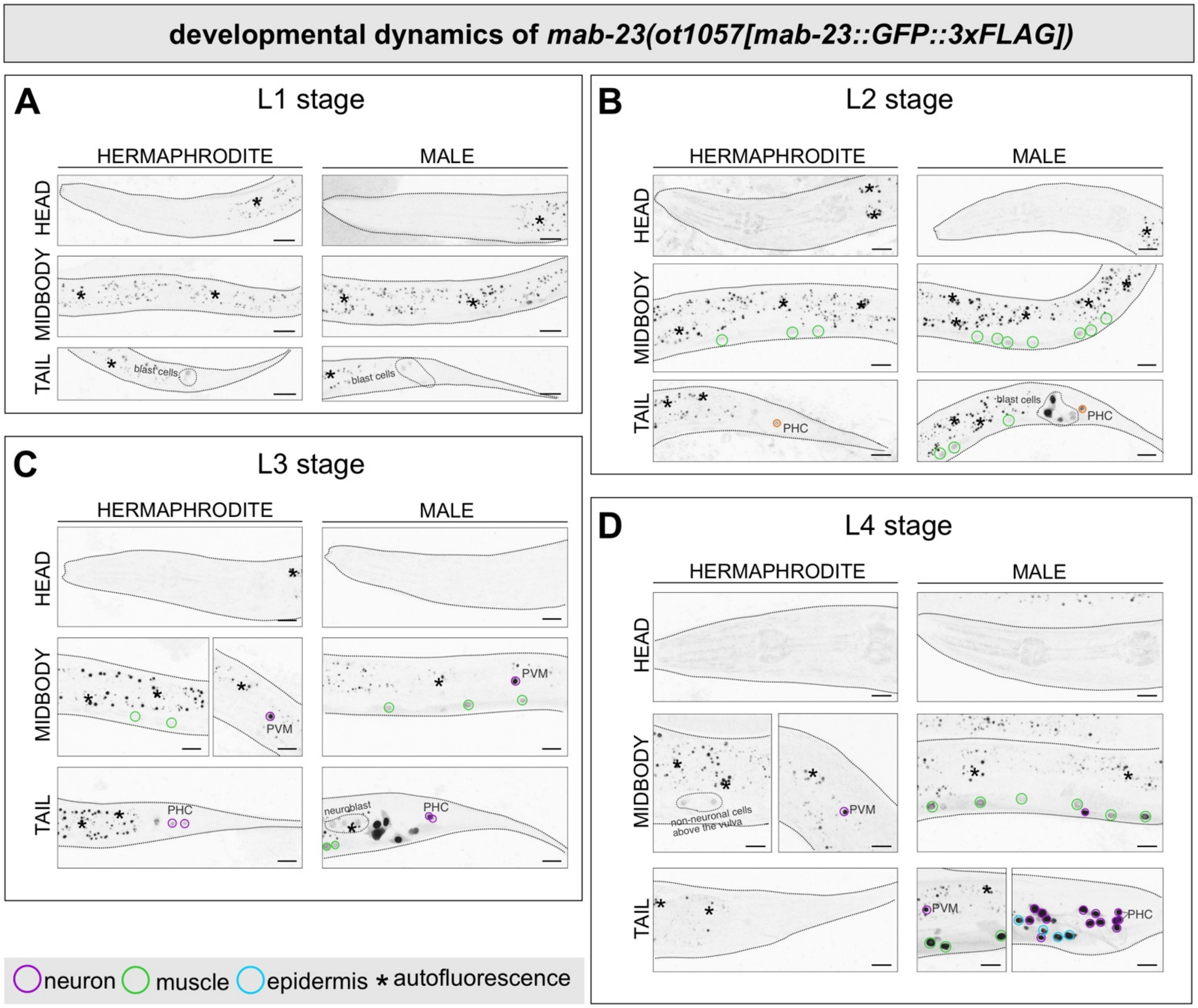

**Figure S4. Temporal dynamics of MAB-23.** *mab-23(ot1057)* reporter allele expression across larval and adult stages in grayscale. (**A-D**) overall expression in L1 (**A**), L2 (**B**), L3 (**C**), and L4 (**D**) stages. Tissue types are outlined in different colors as indicated in the legend on the figure. Refer to **Table 1** and Results for description of expression patterns. Dotted lines indicate the outlines of the animals. Scale bars, 10 µm.

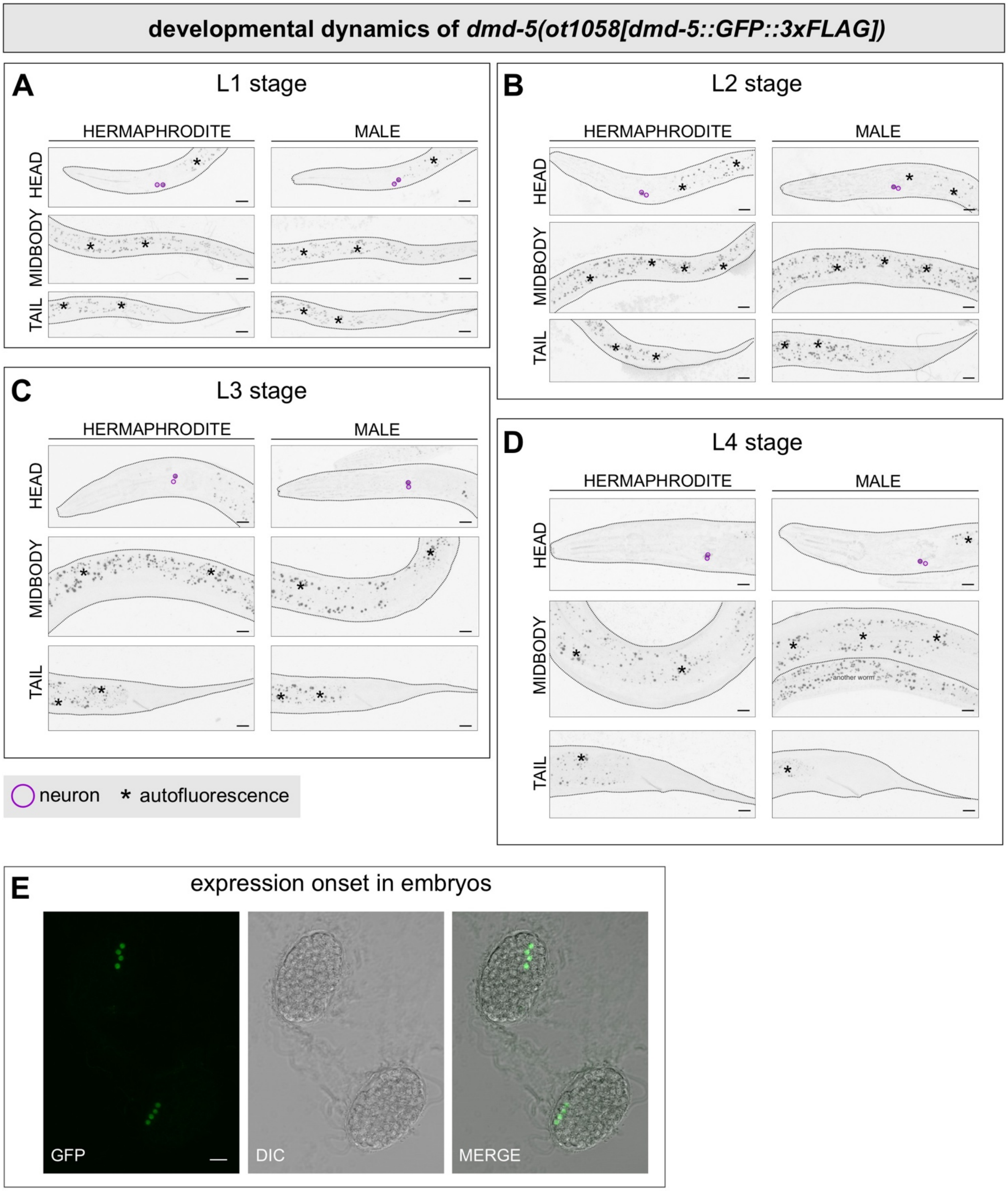

**Figure S5. Temporal dynamics of DMD-5.** *dmd-5(ot1058)* reporter allele expression across larval and adult stages. (**A-D**) overall expression in L1 (**A**), L2 (**B**), L3 (**C**), and L4 (**D**) stages in grayscale. Tissue types are outlined in different colors as indicated in the legend on the figure. (**E**) DMD-5 protein expression is visible from gastrulation in the embryos. Refer to **Table 1** and Results for description of expression patterns. Dotted lines indicate the outlines of the animals. Scale bars, 10 μm.

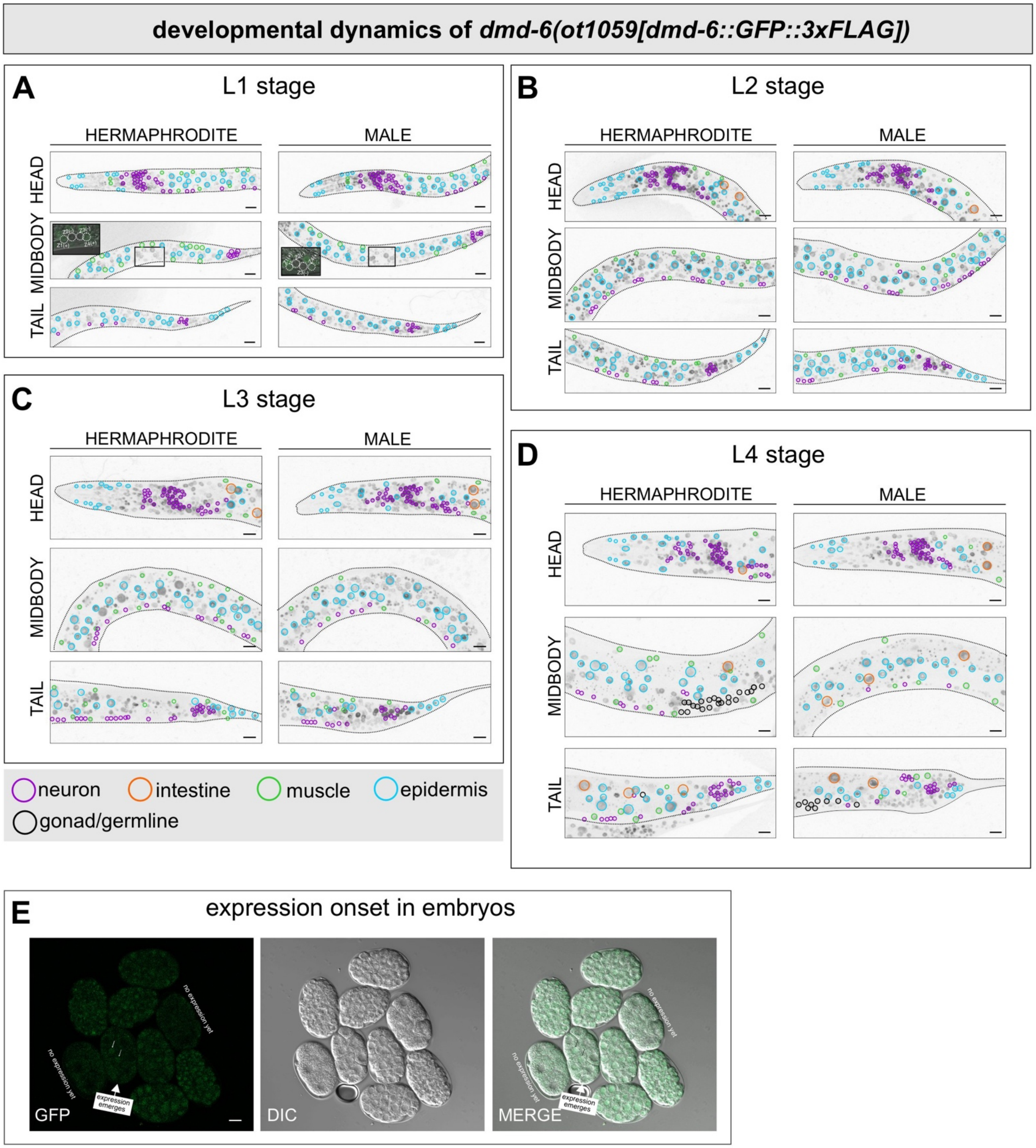

**Figure S6. Temporal dynamics of DMD-6.** *dmd-6(ot1059)* reporter allele expression across larval and adult stages. Images show GFP channels from animals crossed with the ubiquitously expressed marker *mel-28(bq6)* to assess somatic ubiquity. (**A-D**) overall expression in L1 (**A**), L2 (**B**), L3 (**C**), and L4 (**D**) stages in grayscale. Tissue types are outlined in different colors as indicated in the legend on the figure. Cells with ambiguous morphology in densely packed regions are not individually labeled. Precise tissue identification is not required here, as the goal is to determine whether there is broad expression across all somatic cells. (**E**) DMD-6 protein expression is visible from around 8-cell stage in the embryos. Refer to **Table 1** and Results for description of expression patterns. Dotted lines indicate the outlines of the animals. Scale bars, 10 μm.

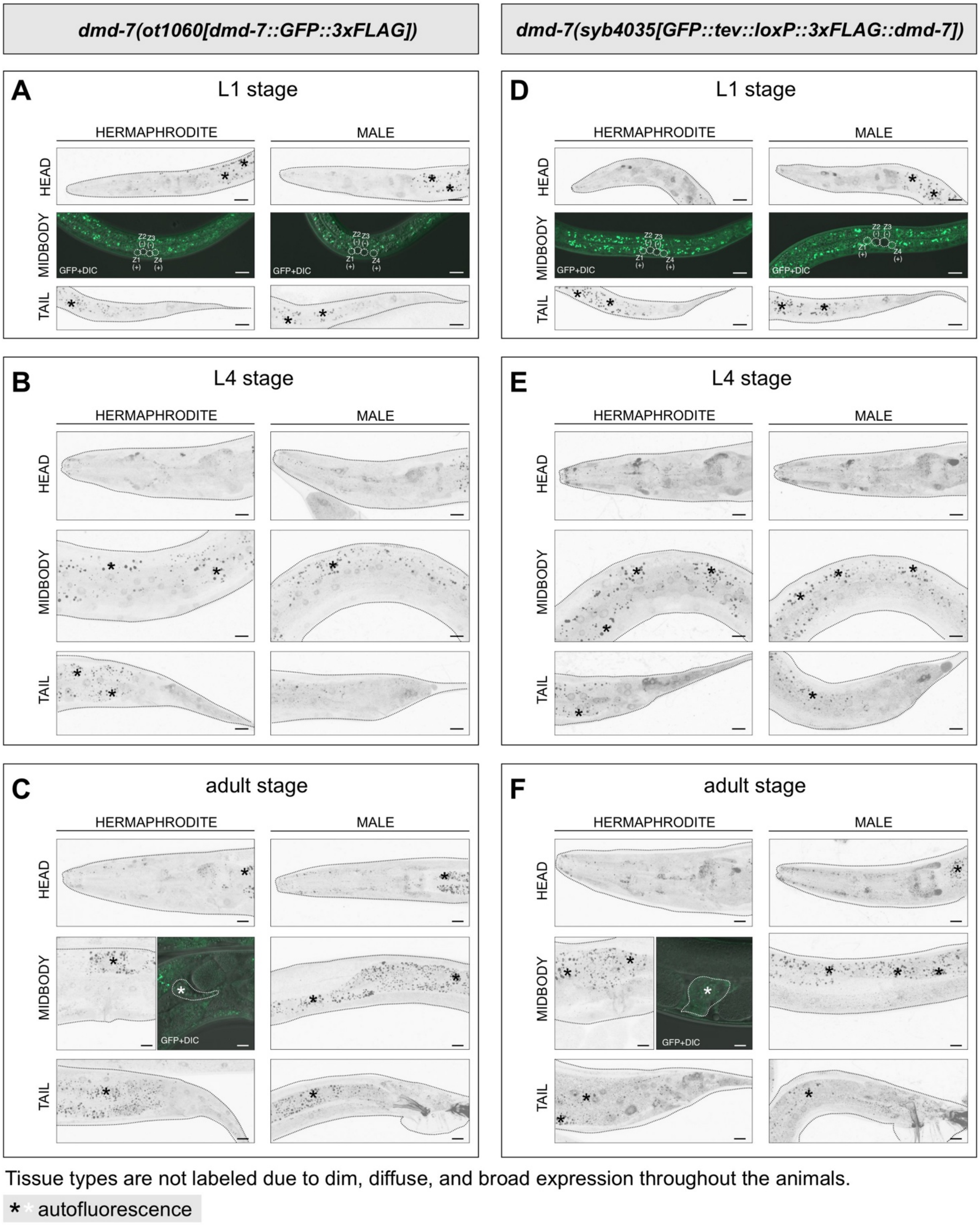

**Figure S7. Temporal dynamic of DMD-7.** Expression of C-terminally tagged reporter allele *dmd-7(ot1060)* and N-terminally tagged reporter allele *dmd-7(syb4035)* across larval and adult stages in grayscale. Both alleles exhibit dim, diffuse, and broad expression in both sexes. (**A-C**) overall expression of *dmd-7(ot1060)* in L1 (**A**), L4 (**B**), and adult (**C**) stages. (**D-F**) overall expression of *dmd-7(syb4035)* in L1 (**D**), L4 (**E**), and adult (**F**) stages. GFP+DIC images in **A** and **D** show the presence of DMD-7 expression in Z1 and Z4, which give rise to somatic gonadal cells, but not Z2 or Z3, which give rise to the germline. GFP+DIC images in **C** and **F** show green signals that are likely autofluorescence in the spermatheca (see also **Fig. S10F**, *him-5* control). Dotted lines indicate the outlines of the animals. Scale bars, 10 μm.

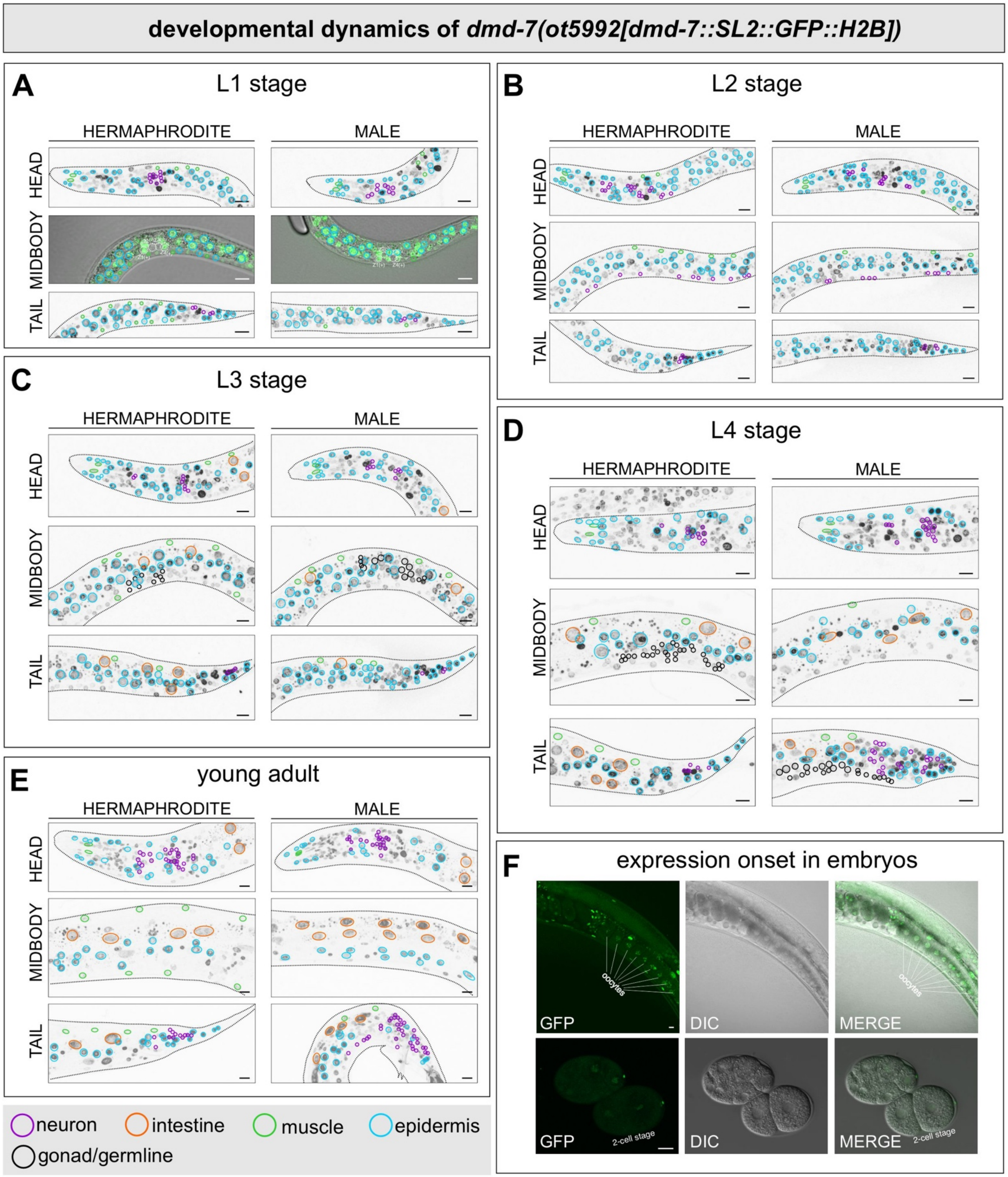

**Figure S8. Expression of an additional *dmd-7* reporter allele in all developmental stages .** *dmd-7(syb5992)* reporter allele expression across larval and adult stages. Images show GFP channels from animals crossed with the ubiquitously expressed marker *mel-28(bq6)* to assess somatic ubiquity. (**A-E**) overall expression in L1 (**A**), L2 (**B**), L3 (**C**), L4 (**D**), and young adult (**E**) stages in grayscale. Tissue types are outlined in different colors as indicated in the legend on the figure. Cells with ambiguous morphology in densely packed regions are not individually labeled. Precise tissue identification is not required here, as the goal is to determine whether there is broad expression across all somatic cells. GFP+DIC images in (**A**) show reporter allele expression in Z1 and Z4, which give rise to somatic gonadal cells, but not Z2 or Z3, which give rise to the germline. This reporter allele is visible in the following neurons in adults: sex-shared neurons expressed in both sexes: ADL, AFD, AIB, AIM, AIN, AIY, ALA, ALN, ASE, ASG, ASH, ASJ, ASK, AVA, AVD, AVE, AVH, AVK, AVM, AWA, AWB, DVA, DVB, DVC, LUA, PDA, PHB, PHC, PLM, PLN, PQR, PVC, PVD, PVN, PVP, PVQ, PVT, PVW, RID, RIP, RIS, SDQ; male-specific neurons: CA1, CA2, CA3, CA5, CA6, CA8, CA9, CP0, CP5, CP6, CP7, CP8, CEM, DVE, DVF, DX1-4, EF1-4, HOB, PGA, R7A, R1B, R2B, R3B, R4B, R5B, R8B, R9B. However, due to its dim and broad expression, we cannot exclude the possibility that it could be in fact ubiquitous in all somatic cells (thus expressed in all neurons). (**F**) Reporter allele expression is visible in oocytes and as early as 2-cell stage embryos. Dotted lines indicate the outlines of the animals. Scale bars, 10 μm.

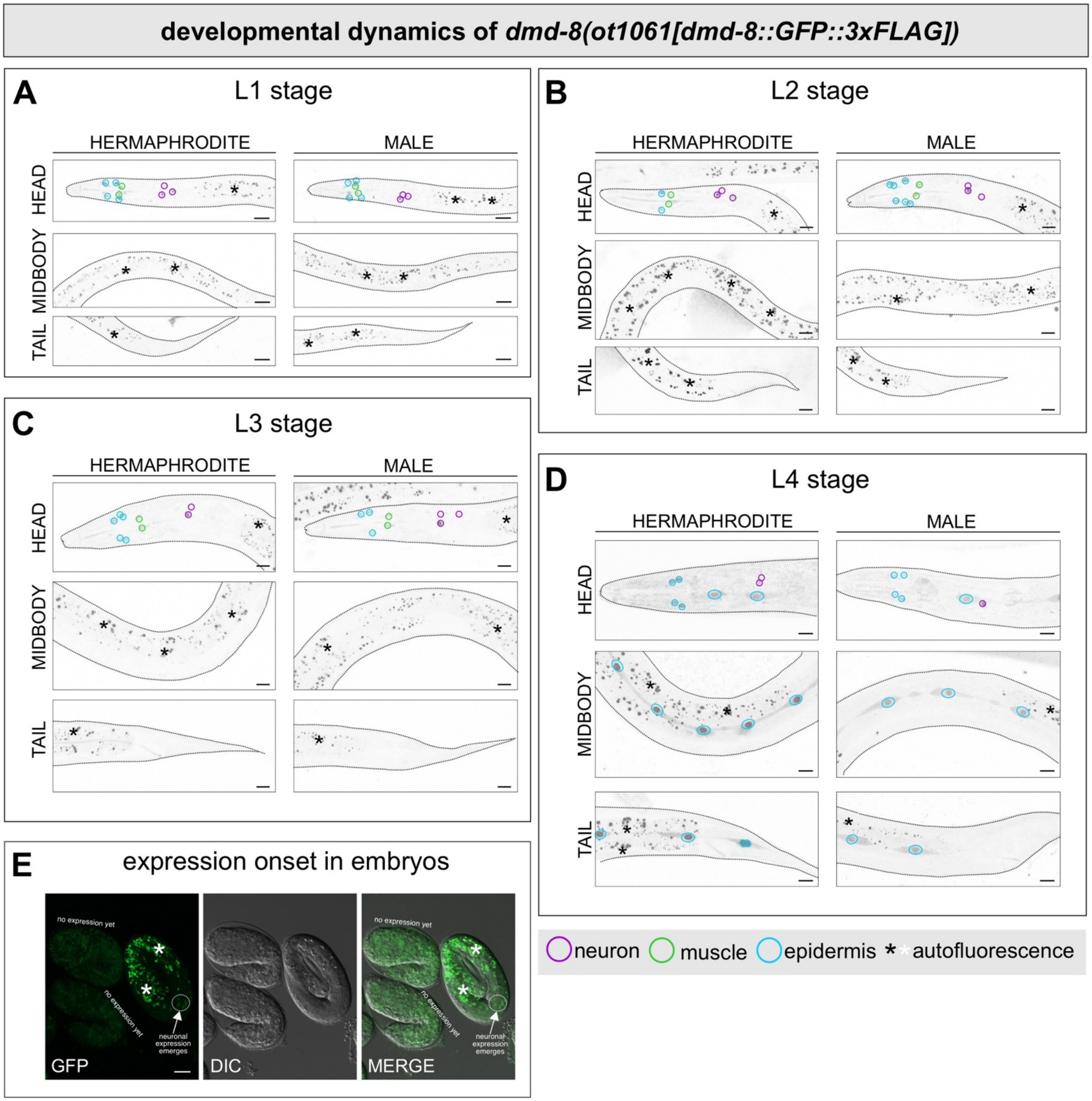

**Figure S9. Temporal dynamics of DMD-8.** *dmd-8(ot1061)* reporter allele expression across larval and adult stages. (**A-D**) overall expression in L1 (**A**), L2 (**B**), L3 (**C**), and L4 (**D**) stages in grayscale. Tissue types are outlined in different colors as indicated in the legend on the figure. (**E**) DMD-8 protein expression is visible from around 3-fold stage in the embryos. Refer to **Table 1** and Results for description of expression patterns. Dotted lines indicate the outlines of the animals. Scale bars, 10 μm.

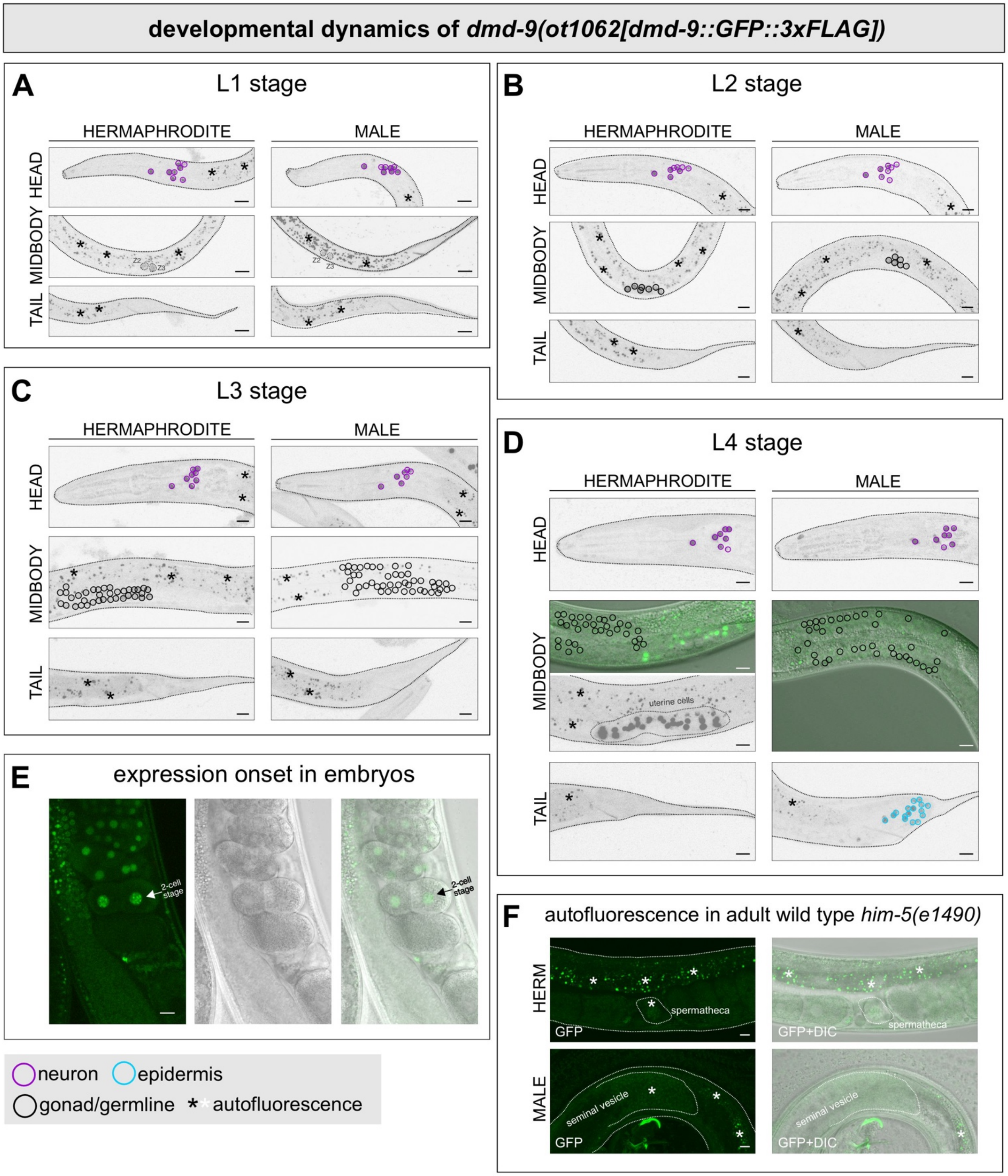

**Figure S10. Temporal dynamics of DMD-9.** *dmd-9(ot1062)* reporter allele expression across larval and adult stages. (**A-D**) overall expression in L1 (**A**), L2 (**B**), L3 (**C**), and L4 (**D**) stages in grayscale. (**E**) DMD-9 protein expression is visible from 2-cell stage in the embryos. (**F**) Adult *him-5(e1490)* control allele imaged using the same settings used for **A-D** and that in **Fig. 4D**. The hermaphrodite spermatheca and male seminal vesicle display autofluorescence. Tissue types are outlined in different colors as indicated in the legend on the figure. Refer to **Table 1** and Results for description of expression patterns. Dotted lines indicate the outlines of the animals. Scale bars, 10 µm.

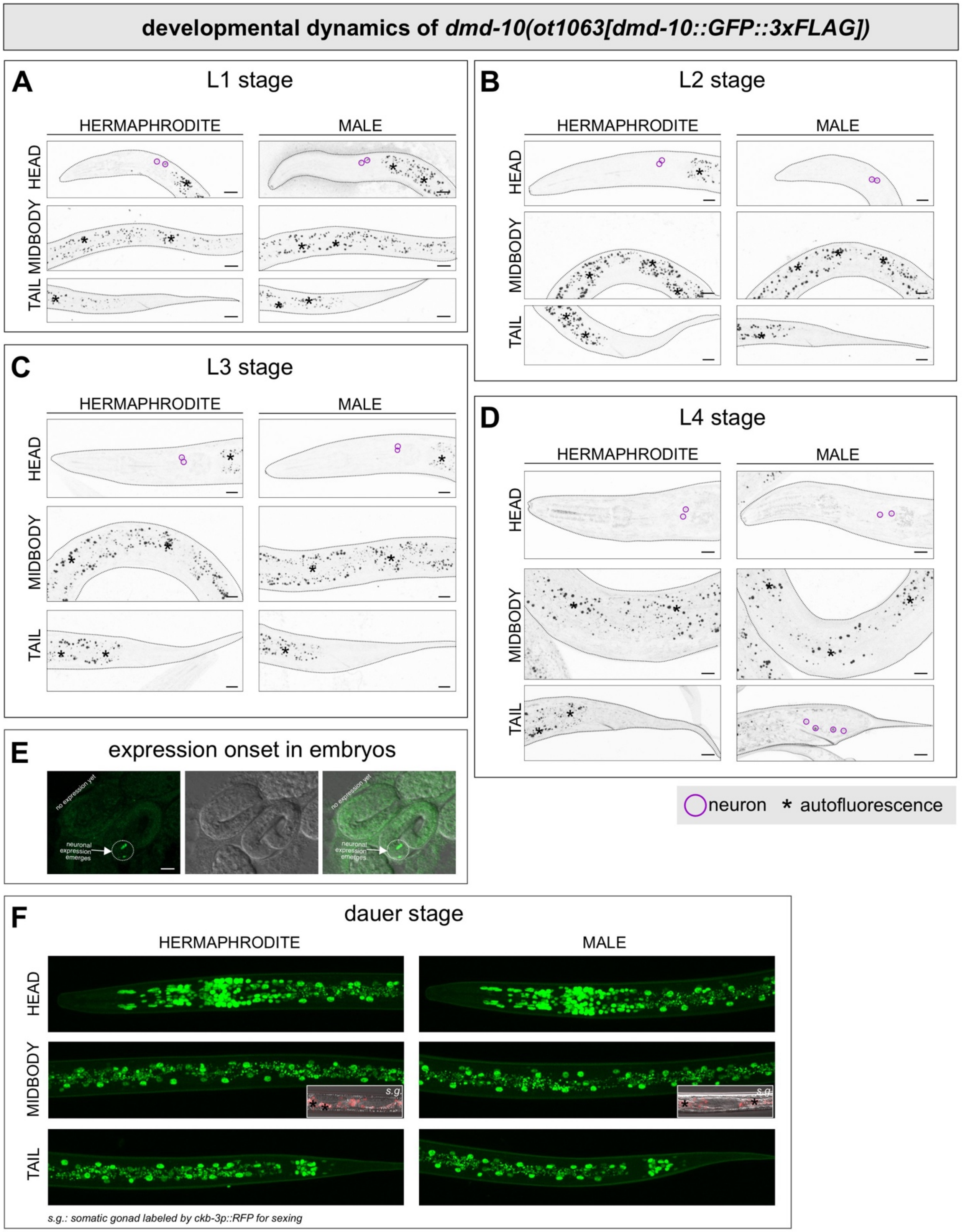

**Figure S11. Temporal dynamics of DMD-10.** *dmd-10(ot1063)* reporter allele expression across larval and adult stages. (**A-D**) overall expression in L1 (**A**), L2 (**B**), L3 (**C**), and L4 (**D**) stages in grayscale. (**E**) DMD-10 protein expression is visible from 3-fold stage in the embryos. Tissue types are outlined in different colors as indicated in the legend on the figure. (**F**) Stress-induced dauer stage animals exhibit broadly upregulated expression in both sexes. Refer to **Table 1** and Results for description of expression patterns. Dotted lines indicate the outlines of the animals. Scale bars, 10 µm.

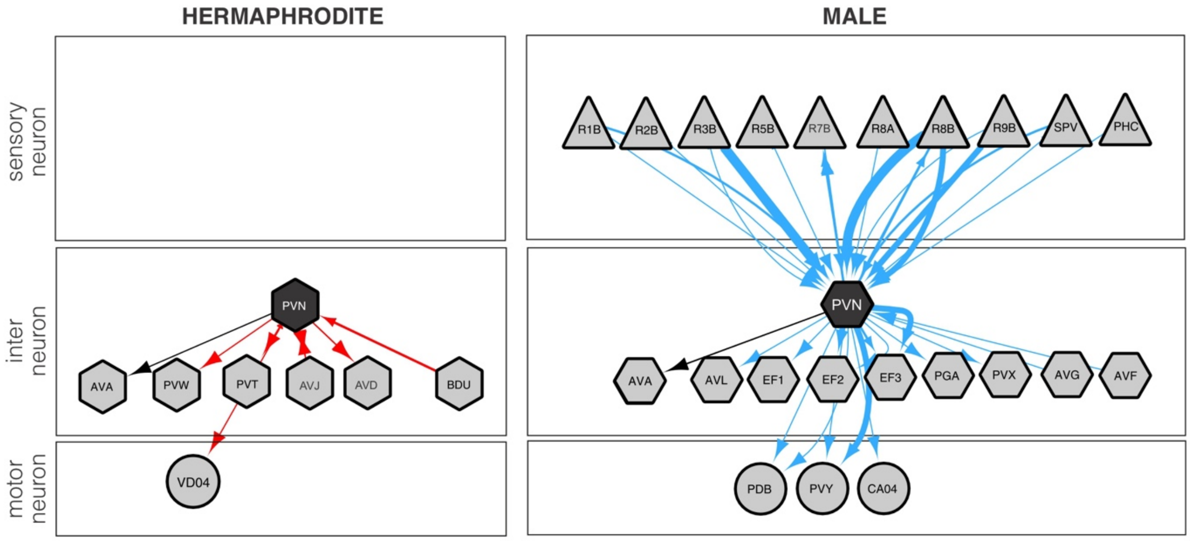

**Figure S12. Chemical synaptic connections of the PVN neuron class.** Thickness of arrows is an indication of the strength of the synaptic connection [46].

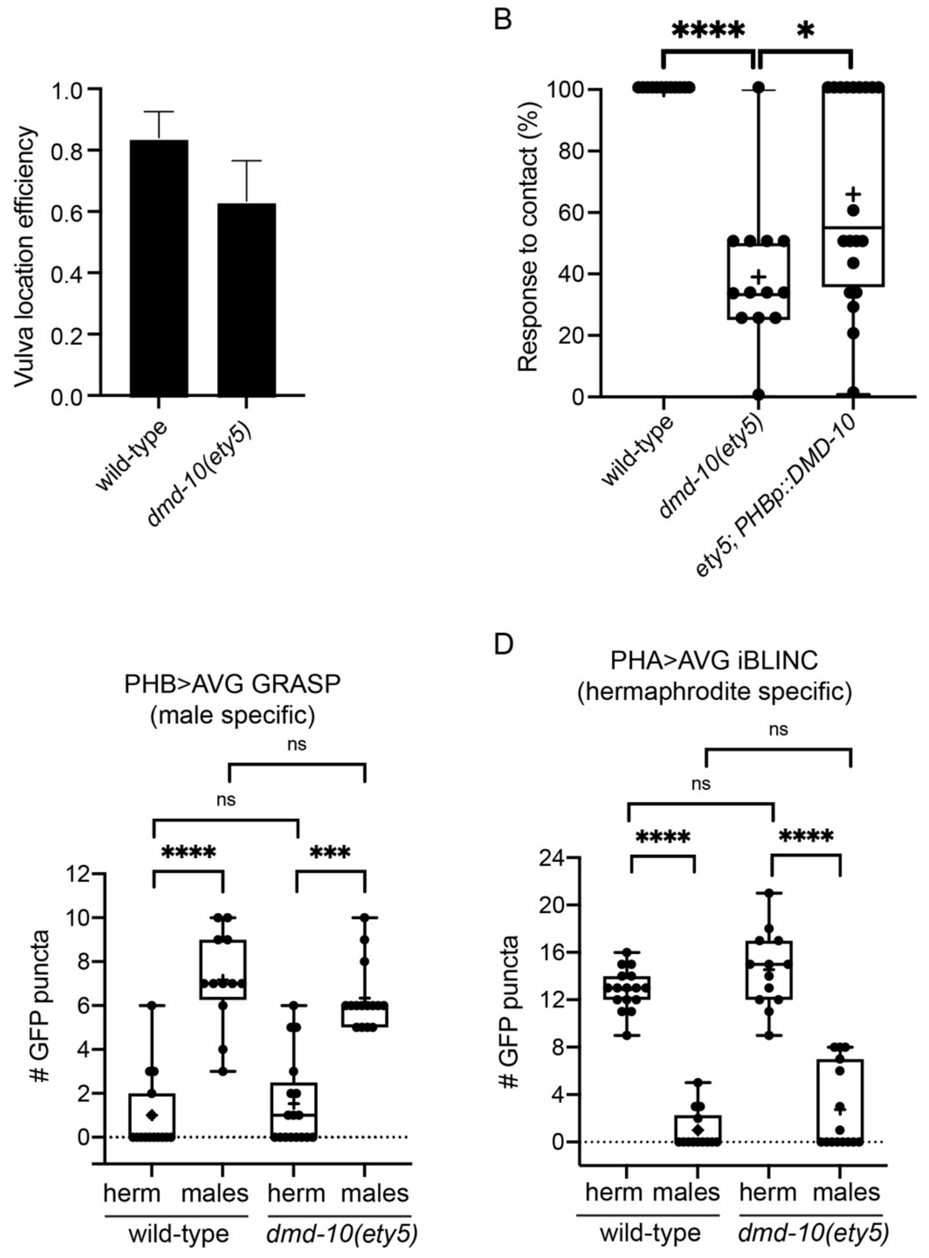

**Figure S13: Functions of *dmd-10* in male-mating behaviors.** *dmd-10* deletion results in a mating defect in males. **(A)** Quantification of vulva location efficiency (see Materials and Methods) in wild type (n=12) vs *dmd-10(ety5)* mutants (n=11). (**B**) Quantification of male response to contact with hermaphrodite during mating in wild type males (n=12), *dmd-10(ety5)* mutants (n=13) and transgenic *ety5* animals carrying a *dmd-10* cDNA rescuing construct specifically in PHB (n=20). **(C-D)** Quantification of synaptic GFP puncta in wild type vs *dmd-10(ety5)* mutants of both sexes. PHB>AVG are trans-synaptically labeled using GRASP [64], PHA>AVG synapses are labeled using iBLINC [65]. PHA>AVG - wild type herm n=16, males n=14, *ety5* herm n=13, *ety5* males n=15. PHB>AVG – wild type herm n=15, males n=12, *ety5* herm n=17, *ety5* males n=15. Statistical tests: A, Mann-Whitney test; B-D: Kruskal-Wallis test followed by Dunn's multiple comparison test.

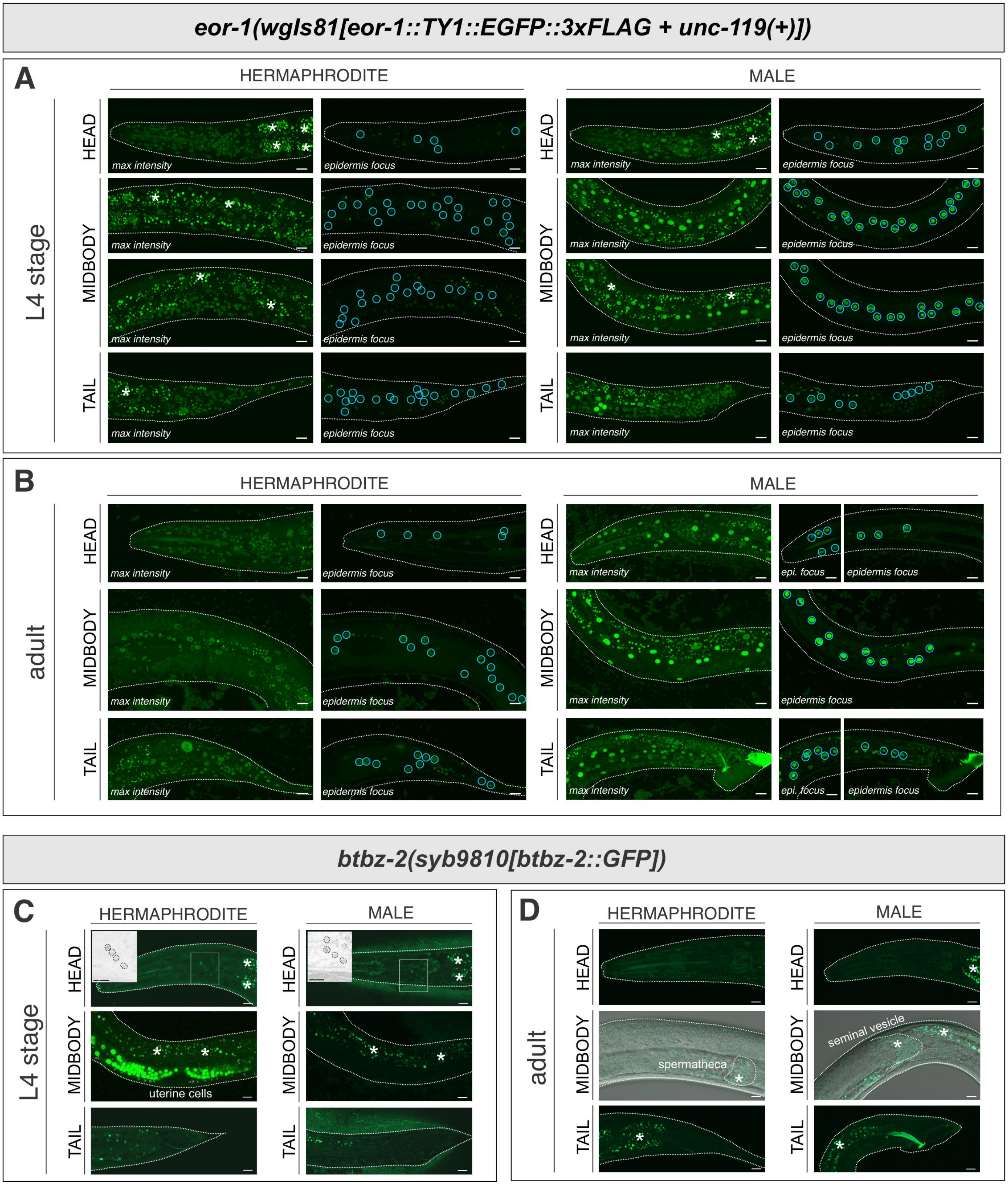

**Figure S14.** ***C. elegans*** **BTB-Zn finger proteins display sexually dimorphic expression patterns.** (**A, B**) *eor-1*(*wgIs81[eor-1::TY1::EGFP::3xFLAG + unc-119(+)])* expression in L4 (**A**) and adult (**B**) stages. Broad expression is observed throughout the animals in both sexes, with higher levels in epidermal (hypodermal) cells in males. For each set of images, maximum-intensity projections are presented on the left, and single focal-plane images focusing on epidermal (hypodermal) cells are presented on the right. Blue circle, epidermal cells. (**C, D**) *btbz-2*(*syb9810[btbz-2::GFP])* expression in L4 (**C**) and adult (**D**) stages. (**C**) BTBZ-2 is expressed dimly in head neurons in both sexes (insets, grayscale images with enhanced contrast and brightness). It is also expressed strongly in hermaphrodite somatic gonad. (**D**) BTBZ-2 continues to express very dimly in head neurons in both sexes and the somatic gonad in the hermaphrodite. For signals in the spermatheca and seminal vesicle, we cannot exclude the possibility that they could be autofluorescence (see also **Fig. S10F**, *him-5* control expression). *, autofluorescence. Dotted lines indicate the outlines of the animals. Scale bars, 10 μm.

| ***C. remanei*** | **Domain structure** (https://smart.embl.de) | **Notes** |
|---|---|---|
| Cre-DMD-4 | 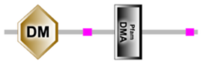 | clear ortholog |
| Cre-DMD-5 | 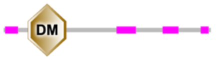 | clear ortholog |
| CRE31496 | 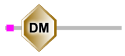 | DMD-5 closest, but NOT the DMD-5 ortholog.<br>Not in immediate vicinity of DMD-5 |
| Cre-MAB-23 | 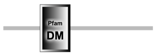 | clear ortholog |
| Cre-MAB-3 | 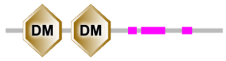 | clear ortholog |
| Cre-DMD-3 | 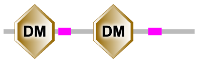 | clear ortholog |
| Cre-DMD-8 | 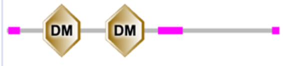 | |
| Cre-DMD-10 + Cre-DMD-11 | | split, as originally in Cel |
| Cre-DMD-6.1 | 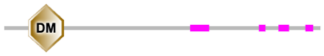 | clear ortholog |
| Cre-DMD-6.2 | same as above | alone on a free-floating contig. Entire nucleotide sequence differs only by 4 nucleotides -> must be the same gene |
| Cre-DMD-7 | 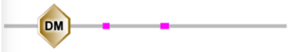 | clear ortholog |
| Cre-DMD-9.1 | 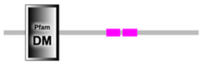 | 9.1/9.2 are almost identical, but real. In close proximity: |
| Cre-DMD-9.2 | 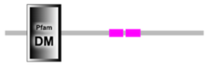 | 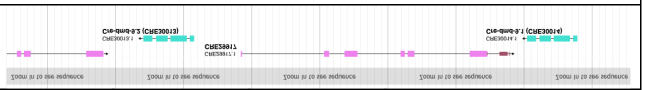 |

| ***O.volvulus*** | **Domain structure** (https://smart.embl.de) | **Notes** |
|---|---|---|
| Ovo-DMD-5 | 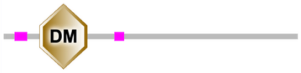 | clear ortholog |
| no DMD-4 | | |
| Ovo-MAB-3 | 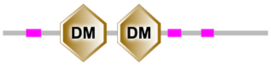 | clear ortholog |
| OVOC6127<br>= MAB-23 | 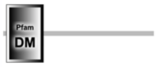 | clear ortholog to MAB-23 |
| Ovo-DMD-3 | 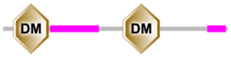 | clear ortholog |
| Ovo-DMD-8 | 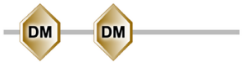 | clear ortholog |
| OVo-DMD-10 + Ovo-DMD-11 | | unfused in Ovo, as originally in Cel |
| Ovo-DMD-7 | 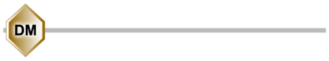 | clear ortholog |
| no DMD-6,9 | | |

| ***P. pacificus*** | **Domain structure** (https://smart.embl.de) | **Notes** |
|---|---|---|
| ppa_stranded_DN18856_c0_g1_i1 (DMD-4) | 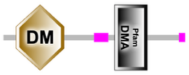 | DMD-4 ortholog |
| ppa_stranded_DN24031_c0_g1_i1 (DMD-5) | 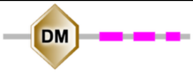 | DMD-5 ortholog |
| PPA37759 ("Ppa-MAB-23") | 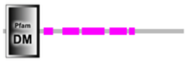 | MAB-23 clear ortholog |
| PPA39038 ("Ppa-DMD-10") | 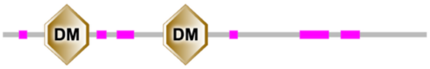 | DMD-10 clear ortholog, properly fused;<br>but also relatively close to DMD-3 (best there is for DMD-3) |
| ppa_stranded_DN20474_c0_g1_i2 ("Ppa-DMD-8.2") | 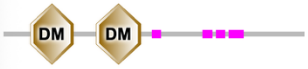 | DMD-8 BRH |
| PPA39495 ("Ppa-DMD-8.1") | 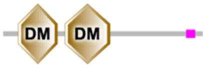 | DMD-8 best hit |
| ppa_stranded_DN15672_c0_g1_i1 ("Ppa-DMD-8.4") | 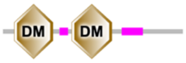 | DMD-8 best hit |
| PPA36867 | 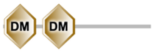 | DMD-8 best hit |
| PPA40793 | 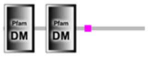 | DMD-8 best hit |
| PPA41401 | 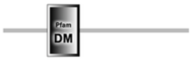 | DMD-8/MAB-3 equal |
| PPA42664 | 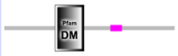 | DMD-8/MAB-3 equal |
| PPA43049 | 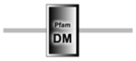 | DMD-8/MAB-3 equal |
| PPA31719 ("Ppa-MAB-3.1") | 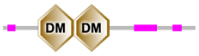 | **MAB-3 best** |
| PPA03790 (" Ppa-MAB-3 = Ppa-MAB-3.2")_ | 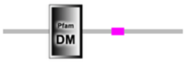 | poor MAB-3 hit |
| PPA06303 | 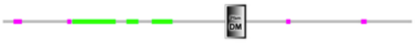 | poor MAB-3 hit |
| PPA29816 | 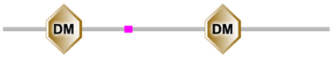 | poor MAB-3 hit |
| ppa_stranded_DN18713_c0_g1_i1 ("Ppa-DMD-6") | 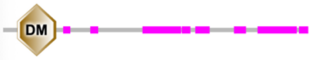 | DMD-6 best hit |
| PPA09037 | 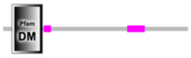 | DMD-7 best hit |
| PPA31139 ("Ppa-DMD-9.1") | 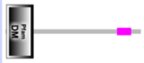 | DMD-7 best hit |
| PPA40600 ("Ppa-DMD-7 ") | 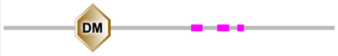 | DMD-7 best hit |
| PPA42389 ("Ppa-DMD-9 = Ppa-DMD-9.2") | 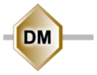 | DMD-7 best hit |
| ppa_stranded_DN17512_c0_g1_i1 | 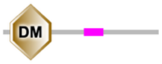 | DMD-9 best hit |

| *P. sambesii* | **Domain structure** (https://smart.embl.de) | **Notes** |
|---|---|---|
| PSAMB.scaffold11507size3306.g34211.t1 | 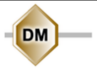  | MAB-3/DMD-3 like |
| PSAMB.scaffold2118size25250.g16466.t1 = PSAMB.scaffold9449size4965.g32452.t1 | 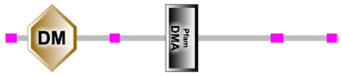  | DMD-4 ortholog |
| PSAMB.scaffold3378size18519.g21250.t1 | 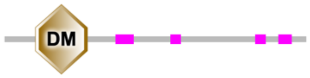  | MAB-3 best |
| PSAMB.scaffold3782size16919.g22525.t1 | 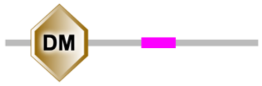  | DMD-10 best |
| PSAMB.scaffold445size50797.g5880.t1 | 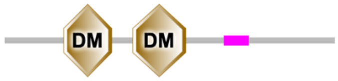  | DMD-3/DMD-10 best |
| PSAMB.scaffold5699size11063.g27140.t1 = PSAMB.scaffold712size42977.g8218.t1 | 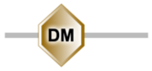  | DMD-5 ortholog |
| PSAMB.scaffold769size41634.g8668.t1 | 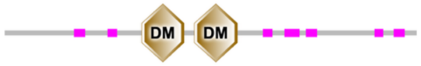  | MAB-3 ortholog |

**Table S1. DMRT sequence analysis in more nematodes**

| | |
|---|---|
| [green] | identity gene expression upregulated |
| [red] | identity gene expression downregulated |
| [grey] | neuron expresses a given DMRT but neurotransmitter identity is not changed |

| | | DM factors expression | | | neurotransmitter pathway genes in wild type | | | | mab-3(-) | | | | mab-23(-) | | | | dmd-3(-) | | | | |
|---|---|---|---|---|---|---|---|---|---|---|---|---|---|---|---|---|---|---|---|---|---|
| Class | Neuron | mab-3 | mab-23 | dmd-3 | eat-4 (glutamatergic) | unc-17 (cholinergic) | unc-25 (GABAergic) | cat-1 (monoaminergic) | eat-4::gfp mab-3(-) | unc-17::gfp mab-3(-) | unc-25::gfp mab-3(-) | cat-1::gfp mab-3(-) | eat-4::gfp mab-23(-) | unc-17::gfp mab-23(-) | unc-25::gfp mab-23(-) | cat-1::gfp mab-23(-) | eat-4::gfp dmd-3(-) | unc-17::gfp dmd-3(-) | unc-25::gfp dmd-3(-) | cat-1::gfp dmd-3(-) | other observations |
| **HEAD** | | | | | | | | | | | | | | | | | | | | | |
| CEM | CEMDL | ✓ | | | | low ++ | | | | | | | | | | | | | | | |
| | CEMDR | ✓ | | | | low ++ | | | | | | | | | | | | | | | |
| | CEMVL | ✓ | | | | low ++ | | | | | | | | | | | | | | | |
| | CEMVR | ✓ | | | | low ++ | | | | | | | | | | | | | | | |
| MCM | MCML | | | | | | | | | | | | | | | | | | | | |
| | MCMR | | | | | | | | | | | | | | | | | | | | |
| **VENTRAL NERVE CORD** | | | | | | | | | | | | | | | | | | | | | |
| CA | CA1 | | | ✓ | very low +/variable | high +++ | | | | | | | | | | | | | | | |
| | CA2 | | | ✓ | very low +/variable | high +++ | | | | | | | | | | | | | | | |
| | CA3 | | | ✓ | very low +/variable | high +++ | | | | | | | | | | | | | | | |
| | CA4 | | | ✓ | very low +/variable | high +++ | | | | | | | | | | | | | | | |
| | CA5 | | ✓ | ✓ | | high +++ | | | | | | | | | | | | | | | NeuroPAL blue lost green gain in dmd-3(-) |
| | CA6 | | ✓ | ✓ | | high +++ | | | | | | | | | | | | | | | NeuroPAL blue lost green gain in dmd-3(-) |
| | CA7 | | ✓ | ✓ | high +++ | low ++ | | | | | | | | | | | | | | | NeuroPAL red blue lost green gain in dmd-3(-) |
| | CA8 | | ✓ | ✓ | | low ++ | | | | | | | | | | | | | | | NeuroPAL green gain in mab-23(-), dmd-3(-) |
| | CA9 | ✓ | ✓ | ✓ | | low ++ | | | | | | | | | | | | | | | NeuroPAL green gain in dmd-3(-) |
| CP | CP0 | | | | low ++ | | | | | | | | | | | | | | | | |
| | CP1 | ✓ | | | | low ++ | | low ++ | | | | | | | | | | | | | |
| | CP2 | ✓ | | | | low ++ | | low ++ | | | | | | | | | | | | | |
| | CP3 | ✓ | | | | low ++ | | low ++ | | | | | | | | | | | | | |
| | CP4 | ✓ | | | | low ++ | | low ++ | | | | | | | | | | | | | |
| | CP5 | | | ✓ | low ++ | | | low ++ | | | | | | | | | | | | | |
| | CP6 | | | ✓ | low ++ | | | low ++ | | | | | | | | | | | | | |
| | CP7 | | ✓ | | | very low + | | | | | | | | | | | | | | | |
| | CP8 | | ✓ | | | very low + | | | | | | | | | | | | | | | |
| | CP9 | ✓ | | ✓ | | no visible expressio | very high ++++ | | | | | | | | | | | | | | |
| **TAIL: DORSALRECTAL GANGLION** | | | | | | | | | | | | | | | | | | | | | |
| DVE | DVE | | | ✓ | | | | | | | | | | | | | | | | | |
| DVF | DVF | | | ✓ | | | | | | | | | | | | | | | | | |
| DX | DX1/2 | | | ✓ | | very low + | | | | | | | | | | | | | | | |
| | DX1/2 | | | ✓ | | very low + | | | | | | | | | | | | | | | |
| EF | EF1/2 | | | ✓ | | | very high ++++ | | | | | | | | | | | | | | |
| | EF1/2 | | | ✓ | | | very high ++++ | | | | | | | | | | | | | | |
| **TAIL: PREANAL GANGLION** | | | | | | | | | | | | | | | | | | | | | |
| HOA | HOA | | | ✓ | very high ++++ | | | high +++ | | | | | | | | | | | | | |
| HOB | HOB | | | | | low ++ | | | | | | | | | | | | | | | |
| PDC | PDC | | ✓ | ✓ | | high +++ | | low ++ | | | | | | | | | | | | | NeuroPAL green gained in dmd-3(-) and mab-23(-) |
| PGA | PGA | ✓ | | ✓ | | low ++ | | very low + | | | | | | | | | | | | | |
| PVV | PVV | | | ✓ | very high ++++ | high +++ | | | | | | | | | | | | | | | NeuroPAL likely dimmer red in dmd-3(-) |
| PVY | PVY | | | ✓ | | very high ++++ | | low ++ | | | | | | | | | | | | | |
| PVX | PVX | | | ✓ | | very high ++++ | | low ++ | | | | | | | | | | | | | |
| PVZ | PVZ | ✓ | | | | low ++ | | | | | | | | | | | | | | | |
| DX | DX3/4 | | | ✓ | | low ++ | | | | | | | | | | | | | | | |
| | DX3/4 | | | ✓ | | low ++ | | | | | | | | | | | | | | | |
| *EF | EF3/4 | | | ✓ | | | very high ++++ | | | | | | | | | | | | | | |
| | EF3/4 | | | ✓ | | | very high ++++ | | | | | | | | | | | | | | |
| **TAIL: LUMBAR GANGLION (RAYS)** | | | | | | | | | | | | | | | | | | | | | |
| RnA | R1AL | | ✓ | ✓ | | very high ++++ | | | | | | | | | | | | | | | NeuroPAL gains red in dmd-3(-) and mab-23(-) |
| | R1AR | | ✓ | ✓ | | very high ++++ | | | | | | | | | | | | | | | NeuroPAL gains red in dmd-3(-) and mab-23(-) |
| | R2AL | | ✓ | ✓ | | very high ++++ | | very low + | | | | | | | | | | | | | NeuroPAL gains red in dmd-3(-) and mab-23(-) |
| | R2AR | | ✓ | ✓ | | very high ++++ | | very low + | | | | | | | | | | | | | NeuroPAL gains red in dmd-3(-) and mab-23(-) |
| | R3AL | ✓ | | | | high +++ | | | neurons invisible in mab-3(-) due to anatomical change | | | | | | | | | | | | |
| | R3AR | ✓ | | | | high +++ | | | | | | | | | | | | | | | |
| | R4AL | ✓ | | | | high +++ | | | | | | | | | | | | | | | |
| | R4AR | ✓ | | | | high +++ | | | | | | | | | | | | | | | |
| | R5AL | | | ✓ | high +++ | | | high +++ | | | | | | | | | | | | | |
| | R5AR | | | ✓ | high +++ | | | high +++ | | | | | | | | | | | | | |
| | R6AL | ✓ | ✓ | ✓ | | very high ++++ | | | neurons invisible in mab-3(-) due to anatomical change | | | | | | | | | | | | NeuroPAL gains red in dmd-3(-) |
| | R6AR | ✓ | ✓ | ✓ | | very high ++++ | | | | | | | | | | | | | | | NeuroPAL gains red in dmd-3(-) |
| | R7AL | | | ✓ | | | | high +++ | | | | | | | | | | | | | NeuroPAL gains red in dmd-3(-) |
| | R7AR | | | ✓ | | | | high +++ | | | | | | | | | | | | | NeuroPAL gains red in dmd-3(-) |
| | R8AL | ✓ | | ✓ | | low ++ | | | | | | | | | | | | | | | |
| | R8AR | ✓ | | ✓ | | low ++ | | | | | | | | | | | | | | | |
| | R9AL | ✓ | | ✓ | low ++ | | | high +++ | | | | | | | | | | | | | NeuroPAL red may be decreased in dmd-3(-) |
| | R9AR | ✓ | | ✓ | low ++ | | | high +++ | | | | | | | | | | | | | NeuroPAL red may be decreased in dmd-3(-) |
| RnB | R1BL | | | | | low ++ | | low ++ | | | | | | | | | | | | | |
| | R1BR | | | | | low ++ | | low ++ | | | | | | | | | | | | | |
| | R2BL | | | | low ++ | | | | | | | | | | | | | | | | |
| | R2BR | | | | low ++ | | | | | | | | | | | | | | | | |
| | R3BL | | | | | | | low ++ | | | | | | | | | | | | | |
| | R3BR | | | | | | | low ++ | | | | | | | | | | | | | |
| | R4BL | | | | | low ++ | | very low + | | | | | | | | | | | | | |
| | R4BR | | | | | low ++ | | very low + | | | | | | | | | | | | | |
| | R5BL | | | | | low ++ | | | | | | | | | | | | | | | |
| | R5BR | | | | | low ++ | | | | | | | | | | | | | | | |
| | R6BL | | | | low ++ | | | | | | | | | | | | | | | | |
| | R6BR | | | | low ++ | | | | | | | | | | | | | | | | |
| | R7BL | | | | | very low +/variable | | | | | | | | | | | | | | | |
| | R7BR | | | | | very low +/variable | | | | | | | | | | | | | | | |
| | R8BL | | | | | low ++ | | | | | | | | | | | | | | | |
| | R8BR | | | | | low ++ | | | | | | | | | | | | | | | |
| | R9BL | | | | | very low +/variable | | low ++ | | | | | | | | | | | | | |
| | R9BR | | | | | very low +/variable | | low ++ | | | | | | | | | | | | | |
| PHD | PHDL | ✓ | | | | low ++ | | | | | | | | | | | | | | | |
| | PHDR | ✓ | | | | low ++ | | | | | | | | | | | | | | | |
| **TAIL: CLOACAL GANGLION** | | | | | | | | | | | | | | | | | | | | | |
| PCA | PCAL | ✓ | | | high +++ | | | | | | | | | | | | | | | | |
| | PCAR | ✓ | | | high +++ | | | | | | | | | | | | | | | | |
| PCB | PCBL | ✓ | ✓ | ✓ | | very high ++++ | | | | | | | | | | | | | | | |
| | PCBR | ✓ | ✓ | ✓ | | very high ++++ | | | | | | | | | | | | | | | |
| PCC | PCCL | ✓ | | | | very high ++++ | | | | | | | | | | | | | | | |
| | PCCR | ✓ | | | | very high ++++ | | | | | | | | | | | | | | | |
| SPC | SPCL | | ✓ | ✓ | | high +++ | | | | | | | | | | | | | | | NeuroPAL blue lost and green gained in dmd-3(-) and mab-23(-) |
| | SPCR | | ✓ | ✓ | | high +++ | | | | | | | | | | | | | | | |
| SPD | SPDL | | | | | | | | | | | | | | | | | | | | |
| | SPDR | | | | | | | | | | | | | | | | | | | | |
| SPV | SPVL | ✓ | | | | high +++ | | | | | | | | | | | | | | | |
| | SPVR | ✓ | | | | high +++ | | | | | | | | | | | | | | | |

* In most cases only one of EF3/4 is visible (Tekieli et al., Genetics, 2021, PMID: 34415309). Here for simplicity we label both neurons as long as we see one expressing a given gene.

### Table S2. DMRT genes broadly regulate neurotransmitter identities in male-specific neurons

**Data S1. Strains used in this study**

**Data S2. Sequences used for CRISPR/Cas9 genome engineering and molecular cloning**
